# GD2-targeting CAR-T cells enhanced by transgenic IL-15 expression are an effective and clinically feasible therapy for glioblastoma

**DOI:** 10.1101/2022.05.01.490250

**Authors:** Tessa Gargett, Lisa M. Ebert, Nga T.H. Truong, Paris M. Kollis, Kristyna Sedivakova, Wenbo Yu, Erica C.F. Yeo, Nicole L. Wittwer, Briony L. Gliddon, Melinda N. Tea, Rebecca Ormsby, Santosh Poonnoose, Jake Nowicki, Orazio Vittorio, David S. Ziegler, Stuart M. Pitson, Michael P. Brown

**Author notes:** Contributed equally (Nga T.H. Truong, Paris M. Kollis, Kristyna Sedivakova). **Correspondence:** Dr Tessa Gargett. **Ethics approval and consent to participate** All donors provided informed consent for the use of their de-identified tissue and blood samples in this study. Use of human glioblastoma patient samples was approved by the Central Adelaide Local Health Network Human Research Ethics Committee (CALHN HREC; approval number R20160727; R20170821). Melanoma patient blood samples for CAR-T manufacturing were obtained under the CARPETS trial protocol (CALHN HREC; approval number R20100524). Samples from DIPG patients were obtained from the Zero Childhood Cancer Initiative (approval number ZCC039), and from the Sydney Children’s Hospital (SCHN HREC; approval 2019/ETH05438). All mouse studies were approved by the University of South Australia Animal Ethics Committee (approval number U44-19). **Consent for publication** All donors provided consent for the publication of their de-identified data in this study. **Availability of data and material** Original datasets (de-identified patient data, scanned microscopy images, flow cytometry FCS files, mouse imaging files, raw impedance values for cytotoxicity assays, mouse clinical records) will be made available by the authors upon reasonable request to the corresponding author (Dr Tessa Gargett, Royal Adelaide Hospital, Adelaide, SA 5000, E). **Authors’ contributions TG:** Conceptualization, investigation, formal analysis, methodology, data curation, writing – original draft, writing – review & editing, funding acquisition. **LME:** Conceptualization, investigation, data curation, funding acquisition, methodology, resources, supervision, writing – review & editing. **NTHT:** Investigation, formal analysis, methodology, data curation. **PMK:** Investigation, formal analysis, methodology, visualization. **KS:** Investigation, formal analysis. **WY:** investigation, methodology. **ECFY:** Investigation. **NLW:** Investigation. **BLG:** Investigation, methodology. **MNT:** Investigation, methodology. **RO:** Resources, data curation **SP:** Resources, data curation **JN:** Resources, data curation. **OV:** Resources. **DSZ:** Resources, data curation. **SMP:** Funding acquisition, resources. **MPB:** Conceptualization, funding acquisition, resources, supervision, writing – review & editing.

## Abstract

**Background:** Aggressive primary brain tumors such as glioblastoma are uniquely challenging to treat. The intracranial location poses barriers to therapy, and the potential for severe toxicity. Effective treatments for primary brain tumors are limited, and 5-year survival rates remain poor. Immune checkpoint inhibitor therapy has transformed treatment of some other cancers but has yet to significantly benefit patients with glioblastoma. Early phase trials of CAR-T cell therapy have demonstrated that this approach is safe and feasible, but with limited evidence of its effectiveness. The choices of appropriate target antigens for CAR-T cell therapy also remain limited.

**Methods:** We profiled an extensive biobank of patients’ biopsy tissues and patient-derived early passage glioma neural stem cell lines for GD2 expression using immunomicroscopy and flow cytometry. We then employed an approved clinical manufacturing process to make CAR-T cells from peripheral blood of glioblastoma and diffuse midline glioma patients and characterized their phenotype and function *in vitro*. Finally, we tested intravenously administered CAR-T cells in an aggressive intracranial xenograft model of glioblastoma and used multicolor flow cytometry, multicolor whole-tissue immunofluorescence and next-generation RNA sequencing to uncover markers associated with effective tumor control.

**Results:** Here we show that the tumor-associated antigen GD2 is highly and consistently expressed in primary glioblastoma tissue removed at surgery. Moreover, despite glioblastoma patients having perturbations in their immune system, highly functional GD2-specific CAR-T cells can be produced from their peripheral T cells using an approved clinical manufacturing process. Finally, after intravenous administration, GD2-CAR-T cells effectively infiltrated the brain and controlled tumor growth in an aggressive orthotopic xenograft model of glioblastoma. Tumor control was further improved using CAR-T cells manufactured with a clinical retroviral vector encoding an IL-15 transgene alongside the GD2-specific CAR. These CAR-T cells achieved a striking 50% complete response rate by bioluminescence imaging in established intracranial tumors. Markers associated with tumor control included those related to T-cell homing, infiltration, and cytotoxicity.

**Conclusions:** Targeting GD2 using a clinically deployed CAR-T therapy has a sound scientific and clinical rationale as a treatment for glioblastoma and other aggressive primary brain tumors.

**What is already known on this topic:** GD2 is a tumor antigen of significant interest for targeting immunotherapy. A single preclinical study has shown the effectiveness of GD2-CAR-T cell therapy in an orthotopic xenograft model of diffuse midline glioma. Similarly, there is one previous preclinical study of GD2-CAR-T therapy in a orthotopic glioblastoma xenograft model but tumor control was achieved only following intracranial injection of CAR-T cells. Given that GD2-CAR-T therapy is already being evaluated clinically for other tumor indications, it is important to establish whether there is an acceptable rationale for its use in brain tumors.

**What this study adds:** This is the first description of a GD2-targeted CAR-T cell therapy that shows antitumor effectiveness in a preclinical model of human glioblastoma following intravenous administration. It is also the first study to investigate the potential effects that the immune profile of glioblastoma patients may have on the feasibility of CAR-T cell manufacturing.

**How this study might affect research, practice, or policy:** The results of this study have led to the initiation of an Australian phase 1 clinical trial program aiming to test GD2-specific CAR-T cells for the treatment of childhood and adult primary brain tumors. The study provides valuable insights into the microenvironmental factors that influence the effectiveness of CAR-T cell therapy for this type of tumor, paving the way for further optimization of CAR-T cell technology for treatment of aggressive primary brain tumors such as glioblastoma.

## Background

Aggressive primary brain tumors have a devastating impact on patients and their families because survival rates are low and treatment options limited. Glioblastoma (GBM), the most lethal of adult gliomas, has a poor 5-year relative survival rate (just 4% at 5 years) which has altered little over 30 years despite use of the multimodal “Stupp” protocol^1^. Recurrence following this treatment is virtually inevitable. The commonest, most lethal form of childhood brain cancer is diffuse intrinsic pontine glioma (DIPG), the major subset of a broader group of diffuse midline gliomas (DMG), which have a midline location. Median survival of children diagnosed with DIPG is approximately 9 months, and overall survival is abysmal with fewer than 1% of patients surviving 5 years. Radiotherapy, the only current standard treatment, increases overall survival by approximately 3 months^2^.

New treatments for these devastating brain tumors represent a high unmet clinical need, and yet these patients have so far not seen much benefit from immune checkpoint inhibitor (ICI) therapy^3^. The other transformative immunotherapy, CAR-T cell therapy, has been widely publicized and successful in treating B-cell malignancies^4,5^. Multiple FDA approvals of CD19-specific CAR-T cell products for patients with relapsed or refractory B-cell acute lymphoblastic leukemia and diffuse large B-cell lymphoma has sparked interest in the application of CAR-T cells to treat a wider range of other cancer types not susceptible to ICIs^6^. CAR-T cell therapy is appealing for glioblastoma because, unlike ICI therapy that requires endogenous tumor-reactive T cells, CAR-T cell therapy relies on an exogenous supply of genetically modified T cells that can be empowered to counter adverse factors generated within the tumor microenvironment. CAR-T cell therapy for GBM is at an early stage of clinical development, with only four reported phase 1 trials (^7-10^ and reviewed in^11^), showing feasibility and safety in targeting the antigens, EGFRVIII, HER2 and IL-13Rα2, with some clinical and biologic evidence of antitumor activity in select patients.

Our chosen target, the glycolipid tumor antigen GD2, is overexpressed in tumors of neuroectodermal origin such as neuroblastoma and has long been a tumor antigen of therapeutic interest^12,13^. The 14g2a monoclonal antibody, from which our CAR single-chain variable fragment (scFv) derives, binds the galactose, N-acetylgalactosamine, and sialic acid regions of the sugar moiety specific to the GD2 molecule^14^. Dinutuximab, the chimeric mAb sharing the same antigen binding domain as the 14g2a scFv, has a standard and FDA-approved role as part of consolidation therapy for high-risk neuroblastoma patients who have achieved at least a partial response to previous multimodal treatment^15^. We and colleagues have investigated third-generation GD2-CAR-T cells as treatment for melanoma, neuroblastoma and other solid cancers^16,17^.

A recent study by Mount et al. has shown high-level GD2 expression in pediatric DMG^18^. Expression of GD2 in glioblastoma has been reported previously in a study of primary cell lines^19^, however expression data direct from patient-derived glioblastoma tissues are limited. In recent publications concerning GD2 as a glioma-associated antigen, investigators have targeted GD2 using mAb therapy^19,20^ or GD2-specific CAR-expressing mesenchymal progenitor cells^21^. In addition, murine GD2-specific CAR-T cells cleared a GD2-expressing murine glioma cell line in a syngeneic, immunocompetent mouse model, but only when used in combination with radiotherapy^22^. In a recent preclinical report of GD2-CAR-T cell therapy in an intracranial model of human GBM, intratumoral administration was effective whereas intravenous administration was ineffective^23^. In addition, this study used a GD2-CAR derived from a monoclonal antibody with no cross-reactivity with mouse GD2, thus preventing the investigators from assessing off-tumor, on-target neurotoxicity.

Here we have validated GD2 as a promising clinical target in adult glioblastoma and confirmed the previously reported findings for DIPG and DMG patients. We have manufactured GD2-specific CAR-T cells with a retrovector in clinical use and performed extensive *in vitro* validation to show that GD2-specific CAR T cells can effectively kill primary DIPG and GBM tumor cells. Importantly, in an *in vivo* model, we demonstrated improved survival following intravenous administration of GD2-CAR-T cells. No on-target, off-tumor neurotoxicity was observed in normal mouse brain, which expresses GD2 and shares the conserved 14g2a epitope with human GD2. Strikingly, the use of an alternative clinical retrovector, which encodes the GD2-specific CAR and a separate IL-15 transgene, produced a CAR-T product that enhanced tumor control.

As an essential proof-of-principle before clinical trials commence, we have also shown that CAR-T cells can be successfully manufactured from PBMCs drawn from GBM or DIPG patients, despite the perturbations that we have observed in the immune compartment of these patients. After GD2-CAR-T cell administration, distinct patterns of tumor infiltration and control were observed in mice, allowing us to uncover tumor microenvironmental factors that may determine the effectiveness of the CAR-T cell therapy. These findings support our clinical investigation of GD2-CAR-T in GBM and DIPG patients (HREC approved, ANZCRT trial registration pending), but also indicate ways to further tailor CAR-based therapies to the unique glioblastoma microenvironment.

## Results

### GD2 is highly expressed in glioblastoma tissues and in glioma neural stem cells derived from patient tissue

We first sought to comprehensively profile GD2 expression in GBM, as published reports of expression levels are limited^19,24,25^, using the same 14g2a antibody clone from which our CAR is derived. To this end, we used an extensive collection of human tumor tissues obtained from the SA Neurological Tumor Bank (a summary of patient material used in this study can be found in **Supplementary Table 1**). From this primary human material, we analyzed both fresh frozen GBM tissue taken at surgical resection and glioma neural stem (GNS) cell lines established as previously described^26^. Importantly, the GNS lines were analyzed at early passage number (<10 for CCB-annotated lines, and <25 for lines obtained as a kind gift from Prof. Bryan Day, QIMRB) and maintained in serum-free media that promote their stem-like state. We found high levels of GD2 expression in all assessed primary patient tissue (n=16) with variable levels of expression in different GBM tissue regions (**Figure 1A-B**). ImageJ was used to independently assess staining intensity and confirm our observation of elevated GD2 staining in all GBM samples compared to tissue identified by the neurosurgeon as adjacent normal brain at the time of resection. To complement this investigation, we performed an analysis of public datasets (**Supplementary Figure 1A-C**), which showed elevated levels of the GD2 synthase enzyme (*B4GALNT1*, responsible for biosynthesis of GD2 and other gangliosides) in GBM, but also its expression in healthy normal brain regions. Nevertheless, direct staining of patient derived tissues indicates low-level expression in non-malignant regions (**Figure 1A**). Early-passage GNS cells were also highly GD2-positive (17 of 20 lines, **Figure 1C-D**). We did not observe GD2 expression on the tissue-culture adapted cell lines, U87 and U251 (data not shown). DIPG has been reported by others to have extremely high GD2 expression^18^. As surgical resection is precluded, we obtained four needle biopsy and four autopsy samples from children with DIPG and confirm enhanced GD2 expression in H3 K27M mutant DIPG (with kind support from the Zero Childhood Cancer initiative, **Supplementary Figure 2**). We also confirmed GD2 expression was maintained when an early passage GNS cell line (CCB-G5c) was established as an intracranial xenograft in NSG mice (**Figure 1E**).

**Figure 1.**
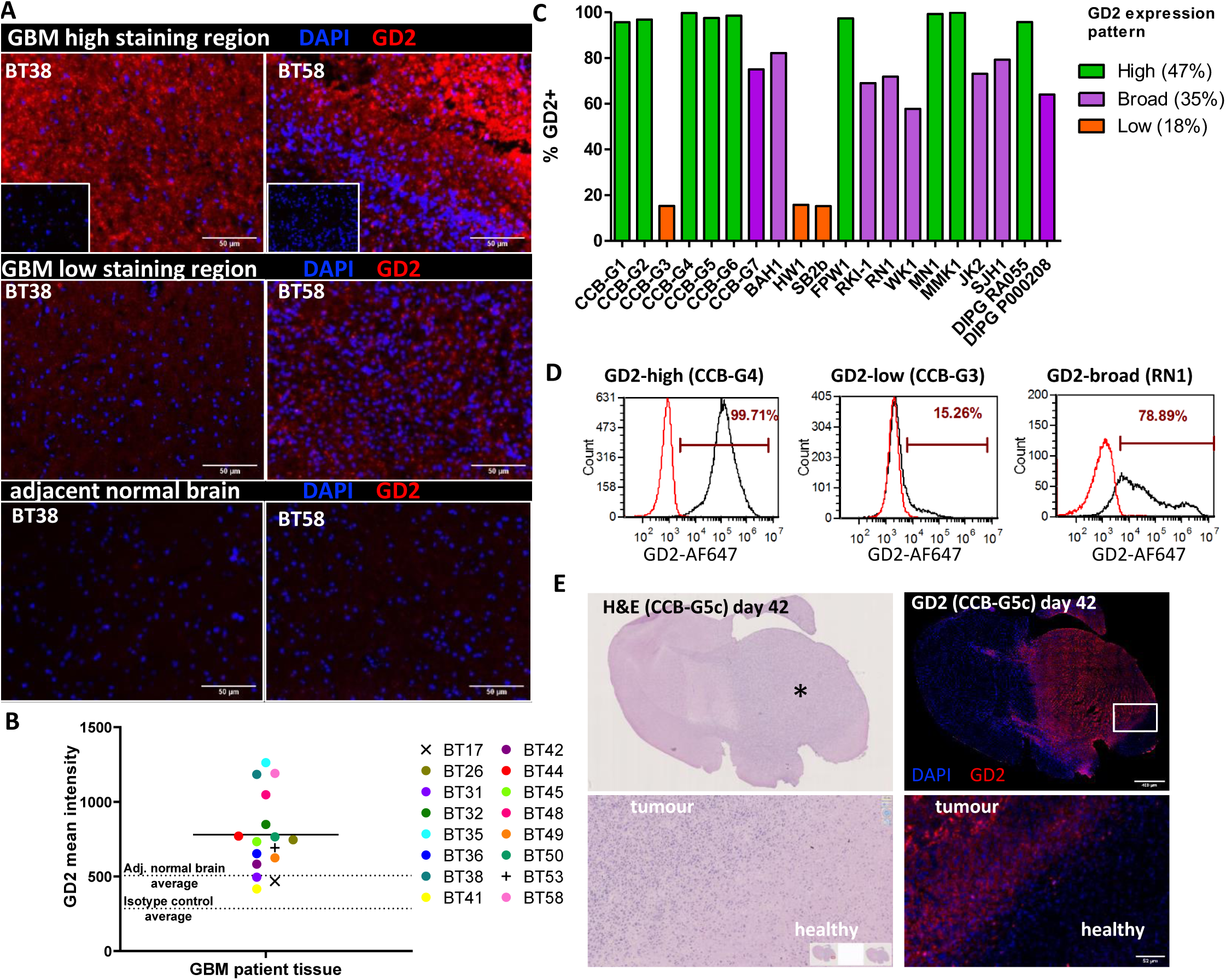
High-level GD2 expression in glioblastoma (GBM) tumor tissues and glioma neural stem (GNS) cell lines, but not in normal brain. **A)** Sections of surgical specimens from GBM (n = 16) or surrounding non-involved brain (n = 4) were stained by immunofluorescence using an anti-GD2 primary antibody (clone 14g2a) and IgG2a isotype control antibody (top row insets). Representative staining for regions of high (top) and low (middle) GD2 expression, and matched adjacent normal brain tissue (bottom). **B)** Summary of GD2 staining intensity measured by imageJ. Dotted lines at y-axis mark average staining intensity for 1) adjacent normal brain tissue (n = 4) removed by neurosurgeon to access the tumor, and 2) the isotype control. **C)** Summary of GD2 expression on GNS cell lines, which were generated from GBM and DIPG patients’ tumors and maintained in culture for < 25 passages. **D)** Representative histograms showing the three distinct GD2 expression patterns observed. Cells were analyzed by flow cytometry for GD2 expression using anti-GD2 primary mAb (clone 14g2a; black histograms) or isotype-matched control antibodies (red histograms). **E)** The CCB-G5C GNS cell line was implanted in the brains of NOD-SCID-gamma-null (NSG) mice via stereotactic intracranial injection. Representative image of hematoxylin and eosin staining (left) and GD2 immunofluorescence (right) of coronal section of mouse brain at time of humane killing because of neurological signs *n* = 10, for full analysis of groups see Figure 4-5. Asterisk marks side of tumor inoculation.

Thus, we found GD2 expression was strongly associated with glioblastoma and DIPG tissues, and GNS cells retained GD2 expression in an aggressive orthotopic mouse model.

### GD2-specific CAR-T cells can be manufactured from glioblastoma patient-derived T cells, but the peripheral immune compartment of these patients is significantly perturbed

Most CAR-T cell therapy occurs in the autologous setting. However, an element often overlooked in preclinical CAR-T cell development is the relative fitness of patient-derived T cells for CAR-T cell manufacturing. In our experience, healthy donor-derived CAR-T cells typically have uniformly successful expansion and potent cytotoxic function, while patient-derived CAR-T cells can have more variable expansion and functional capacity because of patient-specific factors including age, prior treatment, concomitant administration of corticosteroids and disease status. Hence, we undertook an assessment of the suitability of GBM and DIPG patients’ peripheral blood as a starting point for manufacturing, following our established and approved clinical manufacturing protocol for generating third-generation (CD28-OX40-CD3ζ) GD2-CAR-T cells^27^. Donor characteristics are presented in **Supplementary Table 1**, and products were compared to previously manufactured clinical products for melanoma patients on the CARPETS trial (www.anzctr.org.au: ACTRN 12613000198729).

Previously reported findings of lymphopenia, neutrophilia, and a profound decrease in eosinophils in GBM patients^28-30^ were recapitulated in our survey (**Figure 2A** and **Supplementary Figure 3A-D**). Furthermore, compared to melanoma patients, a specific deficit in T cells and an inverted CD4:CD8 ratio, with lower CD4^+^ cells relative to CD8^+^ T cells, was observed both for GBM and DIPG patients in support of a recent report of CD4 T cell bone-marrow sequestration observed with intracranial tumors (**Figure 2B**)^31^. Of note, two melanoma patients with brain metastases had a similarly perturbed CD4:CD8 ratio (open squares). When T-cell immune-phenotyping was performed, significant reductions in the naïve T cell (CCR7^+^, CD62L^+^ and CD45RA^+^) compartment were observed for DIPG and GBM patients compared to melanoma patients (**Figure 2C and Supplementary Figure 3E**). Nevertheless, CAR-T cell manufacturing was achieved for each of six selected patients, with transduction efficiencies and cell expansion yields sufficient to meet our protocol-defined batch release criteria (**Figure 2D**). Cell expansion for several adult GBM patient-derived CAR-T cell products was however slower than that reached for our CARPETS trial melanoma patients. Finally, the CD4:CD8 ratio of CAR-T cell products generally matched that observed in patients’ peripheral blood samples and had a lower proportion of CD4^+^ T cells (**Figure 2E**). Although our culture conditions are optimized for maintaining a central memory phenotype^27^, a smaller proportion of the CAR-T cell product was classified as central memory, and a higher proportion had a heterogenous TEMRA phenotype compared to melanoma patient-derived CAR-T cells (**Figure 2F** and **Supplementary Figure 3E**).

**Figure 2.**
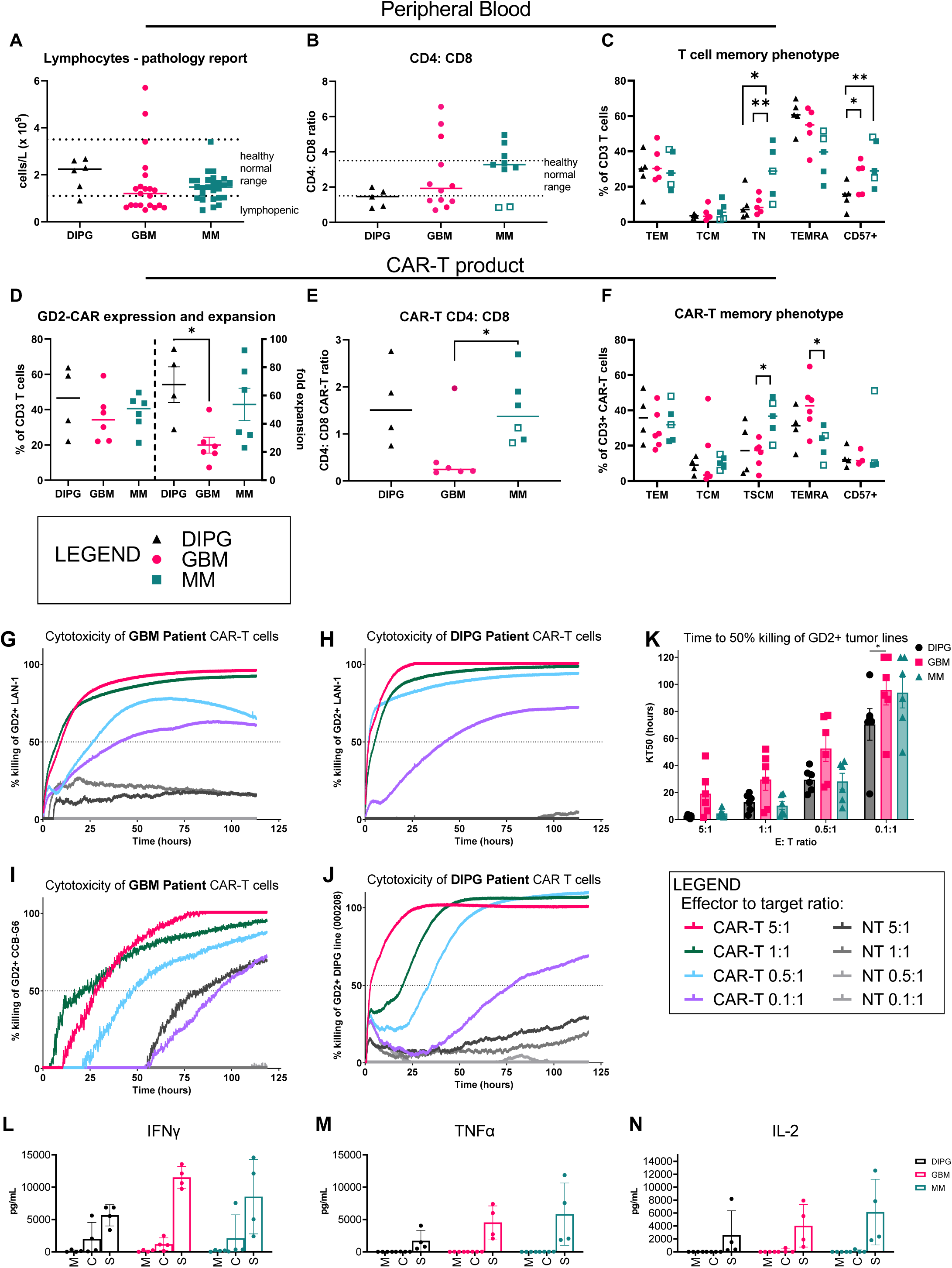
GD2-specific CAR-T cells can be manufactured from peripheral blood of glioblastoma (GBM) and diffuse intrinsic pontine glioma (DIPG) patients and have a distinct phenotype. Immune phenotype of GBM, DIPG, and metastatic melanoma (MM) patient-derived CAR-T cells as determined by multicolor flow cytometry. **A)** Peripheral blood lymphocyte counts obtained at time of blood collection for CAR-T cell manufacture. **B)** Ratio of peripheral blood lymphocyte CD4^+^ and CD8^+^ T cells for each tumor type. Pathology service-defined healthy normal range is marked where available. **C)** Proportions of various memory subsets within the peripheral T-cell population: effector memory (CD45RA^-^ CCR7^-^ CD62L^-^); central memory (CD45RA^-^ CCR7^+^ CD62L^+^); naïve (CD45RA^+^ CCR7^+^ CD62L^+^); TEMRA (CD45RA^+^ CCR7^-^ CD62L^+/-^). **D)** Relative expression and expansion of GD2-CAR-T cells *in vitro* for each tumor type. **E)** Ratio of CD4^+^ and CD8^+^ CAR-T cells. **F)** Proportions of memory subsets within CAR-T cells, defined as for **C)**. Cytotoxicity against GD2-expressing tumor cell lines of neuroblastoma (LAN-1), GBM (CCB-G6) and DIPG (000208) as determined by a real-time cell adhesion-based assay. 0 hours is the time from CAR-T cell addition, after tumor cultures were established overnight. CAR-T and non-transduced control T cells (NT) were added to adherent tumor cells at a range of effector to target ratios. CAR-T cells derived from **g)** GBM (BT29) and **H)** DIPG (DIPG1) patients were assayed against the LAN-1 neuroblastoma target cell line used for batch release testing in our clinical trials. CAR-T cells derived from **I)** GBM (BT29) and **J)** DIPG (DIPG1) patients were assayed against the matched glioma neural stem cell line (CCB-G6) and the unmatched DIPG cell line (000208), respectively. Representative data from one patient are shown; *n* = 4 patient samples. **K)** Summary data of all patient product cytotoxicity assays showing the time in hours to reach 50% killing of targets (KT50). Production of **L)** IFN-gamma **M)** TNF-alpha **N)** IL-2 from CAR-T cell products from DIPG, GBM or metastatic melanoma (MM) patients were cultured with media only (M), media plus IL-7 and IL-15 homeostatic proliferative cytokines (C), or media and plate-bound 1A7 antibody (2µg/mL) for CAR stimulation for 72 hours (S). Other cytokines, and cytokines from peripheral T cells directly isolated from patients are shown in **Supplementary Figure 5**. Supernatants were analyzed by Legendplex Cytometric Bead Array Human Essential Immune Response panel; *n* = 3 patient samples.

To assess the function of CAR-T cells in vitro, we co-cultured a range of GD2^+^ tumor cell lines with CAR-T cells in a real-time impedance-based cytotoxicity assay over five days. CAR-T cells manufactured from GBM patients (**Figure 2G and I)** and DIPG patients (**Figure 2H and J**) effectively killed GD2^+^ cells of the neuroblastoma line, LAN-1, and cells of both early-passage GNS and DIPG lines. We use the level of LAN-1 cytotoxicity as a CAR-T cell manufacturing batch release criterion. Importantly, the GBM line was efficiently killed by CAR-T cells manufactured from the same GBM patient donor. Killing was also achieved at low effector: target ratios of 1 CAR-T cell to 10 tumor cells, indicating serial killing or a bystander killing effect, and DIPG-derived CAR-T cells were particularly potent, with the shortest KT50 at this lowest ratio (**Figure 2K**).

Cytokine production from stimulated CAR-T cells indicated they secreted multiple effector cytokines including IFNγ, TNFα and IL-2, and did so at a comparable level to CAR-T cells manufactured from our melanoma patients (**Figure 2L-N**). Immune-suppressive cytokines IL-10 and TGFβ were undetectable (not shown), however inflammatory cytokines such as IL-17A, IL-6 and IL-8 were also significantly produced, most markedly from the DIPG-patient derived CAR-T cells and this matched what was seen for peripheral T cells sorted directly from these patients’ blood (**Supplementary Figure 4**).

Therefore, these data show that although GBM and DIPG patient-derived CAR-T cells had a distinct phenotype, they performed well *in vitro* in functional assays of cytotoxicity and cyotokine secretion.

### GD2-specific CAR-T cells control orthotopic xenografts of glioblastoma

To determine CAR-T activity *in vivo* we employed an orthotopic xenograft model. In this model NSG mice, receive an intracranial injection of an early-passage patient-derived GNS cell line carrying a luciferase reporter gene *via* stereotactic delivery to the right brain hemisphere (**Figure 3**). The CCB-G5c line was selected for its rapid growth kinetics, which cause neurological signs such as head-tilt and altered gait between days 35 and 42. These clinical signs are used, in part, as the ethical endpoints for humane killing. We sought to compare treatment with healthy donor-derived CAR-T cells with GBM-patient derived CAR-T cells to determine whether differences in their *in vivo* fitness and tumor control were evident.

**Figure 3.**
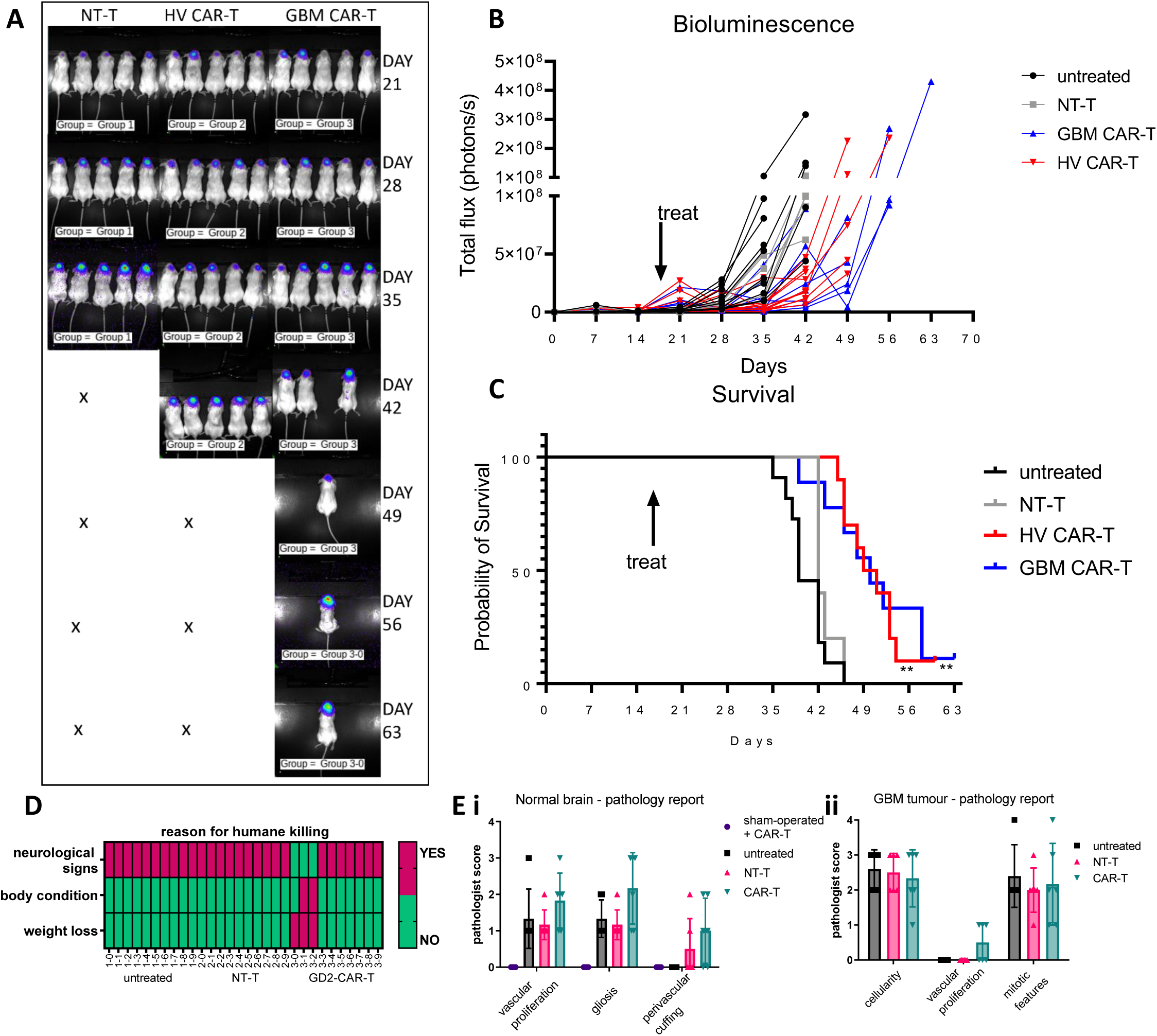
Third-generation GD2-CAR T cells control orthotopic GBM xenografts but do not adversely affect the normal mouse brain. Mice received 2 × 10^5^ CCB-G5c cells by stereotactic intracranial injection on Day 1. On Day 17, mice were given single intravenous injections of saline (untreated), 1.5 × 10^6^ non-transduced control T cells from a healthy donor (Non-transduced T), 1.5 × 10^6^ healthy donor-derived GD2-CAR-T cells (HV CAR-T), or 1.5 × 10^6^ GBM patient-derived GD2-CAR-T cells (GBM CAR-T, from donor BT11 or donor BT48, unmatched to the xenograft). *n* = 8-10/group. **A)** Representative bioluminescence imaging (BLI) for NSG mice with intracranial GBM xenografts. **B)** BLI data for all mice. **C)** Kaplan-Meir survival curves and statistics for mice. Statistics shown are from individual Gehan-Breslow-Wilcoxon tests comparing each curve to the untreated curve. **D)** Clinical signs resulting in humane killing according to pre-defined criteria (see Methods). Neurological signs are one or more of head-tilt, circling, and loss of balance. Mice humanely killed for neurological signs also routinely displayed weight-loss, reluctance to move and ruffled coats, however these alone did not reach a severity score requiring euthanasia. **E)** Independent histopathology scoring of brain sections from humanely killed mice of the **i)** remaining normal brain tissue and **ii)** GBM tumor. Abnormal features were graded as: 0 = none; 1 = minimal; 2 = moderate; 3 = severe. Cellularity was graded as: 1 = low; 2 = moderate; 3 = dense. Mitotic features were reported as number per field of view.

Bioluminescence imaging revealed a delayed increase in luminescence signals for mice treated with CAR-T cells from healthy donors (HV CAR-T) or GBM patients (GBM CAR-T) compared to controls: untreated mice or mice treated with non-transduced T cells (NT-T) (**Figure 3A-B**). Survival was significantly longer in CAR-T treated mice (mean 52 days, HV CAR-T and 53 days, GBM CAR-T) compared to untreated or NT-T treated mice (each mean, 42 days) (**Figure 3C**). CAR-T treated mice were also less likely to display neurological signs and were more frequently euthanized because of poor body condition or weight loss or both, which were the other ethical endpoint criteria for the study (**Figure 3D**). Here, we note that these constitutional signs may be disease- or treatment-related because NSG mice are susceptible to graft versus host disease (GvHD) with GvHD onset reported at 4-6 weeks in other studies^18^. Importantly, no CAR-T cell treated mice were killed for neurological signs before the control tumor-bearing mice consistent with a lack of GD2-CAR-T-cell-mediated neurotoxicity, which had been observed in other studies of GD2-CAR-T cell therapy as early as day 7 post-infusion^18,32^. We employed independent clinical and veterinary pathologists to evaluate histopathologic samples from the xenograft tumors and normal mouse brain tissue, respectively (**Figure 3E**) with no abnormal findings reported for normal brain tissues. Sham-operated mice, which received an intracranial injection of saline, were also treated with CAR-T cells to assess whether neurotoxicity was evident in the absence of tumor and no pathologic signs were detected (**Figure 3Ei)**

Accordingly, we could show that both healthy and GBM patient-derived GD2-CAR-T cells slowed glioblastoma growth and did not contribute to on-target, off-tumor neurotoxicity, the treatment was not curative in this model. We hypothesized that one, or more of the following, were the likely cause for the tumor escaping CAR-T control: 1) insufficient CAR-T homing to and infiltration of the tumor, 2) insufficient CAR-T cell function because of intrinsic T-cell defects or extrinsic tumor microenvironmental factors, or 3) rebound growth of GD2-negative tumor. To determine which factors contributed to immune escape, we next undertook a detailed analysis of the brain tissues using multicolor immunofluorescence and high-parameter flow cytometry.

### Distinct patterns of CAR-T cell tumor infiltration and CD31^+^ vessel formation at study endpoints correlate with survival

Immunofluorescence staining for GD2 showed that in most (9 of 11) CAR-T treated mice, GD2 was still abundantly expressed in the tumor, ruling out tumor antigen loss as a reason for the majority immune escape **(Figure 4**). Nevertheless, the single longest-surviving mouse, which had a notable number of human CD3^+^ tumor-infiltrating T cells **(Figure 4A; iv)**, also had the lowest level of tumor GD2 expression but with a strong tumor bioluminescence signal and high-level human-specific GFAP (hGFAP) staining (**Figure 6**), indicating that in this case, GD2-negative or GD2-low tumor cells had resulted in tumor regrowth (**Figure 4A-C)**. Analysis of human CD3^+^-stained tumor-infiltrating T cells revealed marked variations in the extent of T-cell infiltration at study endpoints. T-cell infiltration in treated mice was graded as low/negative, intermediate, or high (T-cell inflamed) (**Figure 4A-C**). Of note, high levels of CD3 staining correlated with the longest survival (R squared 0.6375, p=0.0004).

**Figure 4.**
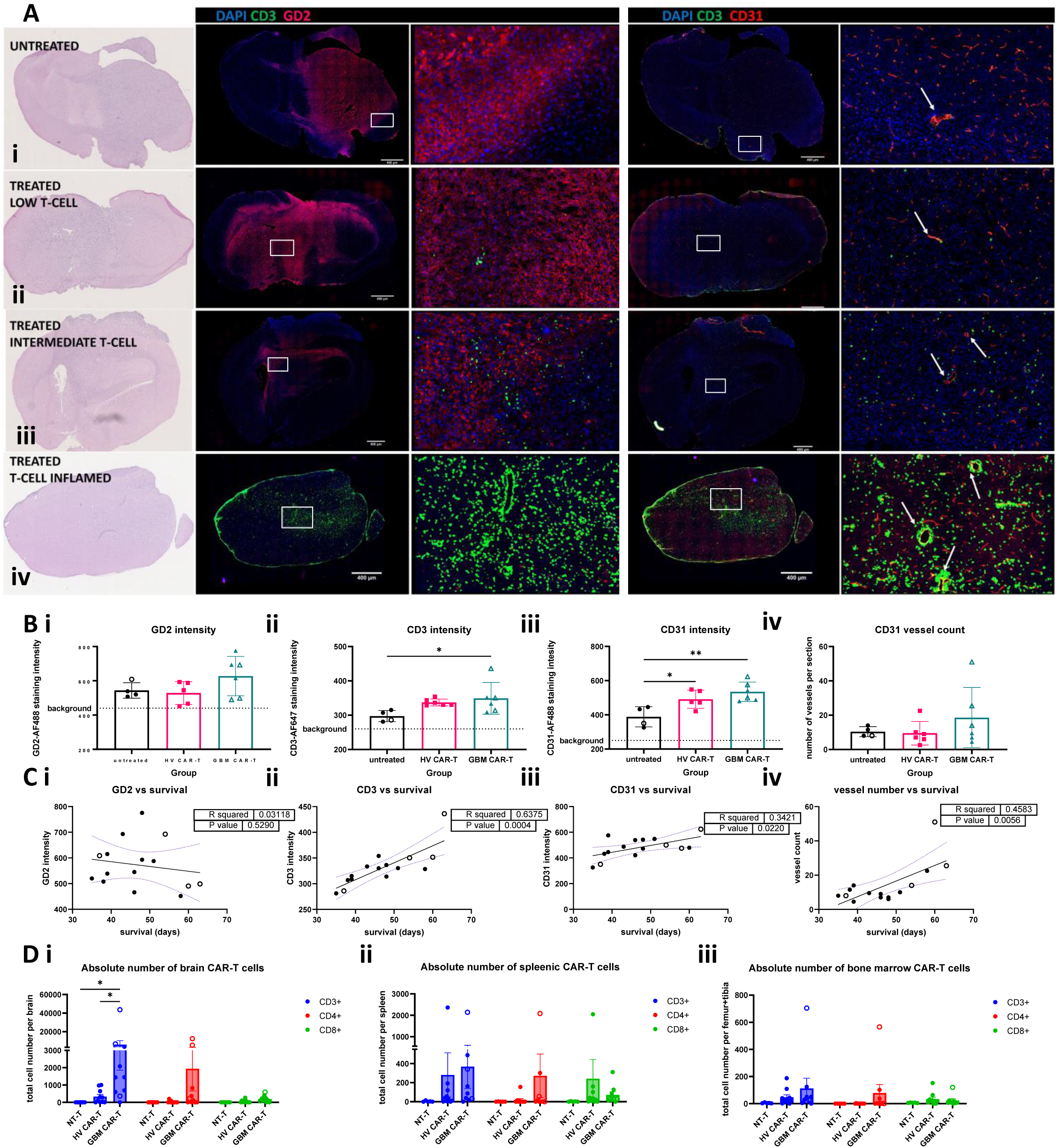
T-cell infiltration as shown in whole brain sections and dissociated tissue samples. At the time of humane killing, mouse brains were bisected, and half was reserved for immunofluorescence (IF) staining of whole tissue, and half was dissociated for flow cytometric analysis alongside spleen and bone marrow. **A)** Whole brain sections (mid-coronal plane where possible) were assessed by H&E staining (far left column), and immunofluorescence using anti-GD2-AF488 (clone 14g2a) and anti-human CD3 AF647 (columns 2 and 3) or anti-human CD3 AF647 and anti-mouse CD31-AF488 (columns 4 and 5) antibodies. White boxes on the whole brain sections indicate regions of interest shown at higher magnification (40x) to the right. White arrows indicate large vessels, identified by the presence of a black lumen and distinct from microvessels. Representative images from i) an untreated mouse, and three GBM CAR-T treated mice (ii-iv) have been chosen to show the range of staining for each molecule. *n* = 6/group. **B)** ImageJ analysis of the staining intensity for GD2, CD3 and CD31 (i-iii), and enumeration of the number of large CD31^+^ vessels (iv). The individual mice represented in the IF microscopy images are shown with open symbols. **C)** Linear regression analysis of staining intensity (i-iii) or large vessel number (iv) vs survival time of mice. **D)** Flow cytometric analysis of dissociated i) brain, ii) spleen and iii) bone marrow tissues using Trucount beads to determine absolute numbers, and the anti-idotypic 1A7 antibody to identify CAR-T cells. *n* = 8-10/group. See **Supplementary figure 6** for representative dot plots and statistical analysis of the correlation between absolute cell numbers and survival.

Staining for CD31 highlighted the abundance of micro-vessels within the tumor. It also showed T cells surrounding larger structured vessels, and there was a significant correlation between numbers of large vessels and survival (R squared 0.4583, p=0.0056, **Figure 4A-C**). Large vessels within the GBM xenograft were not observed in untreated mice, suggesting this was not a tumor-intrinsic feature, but a feature resulting from CAR-T cell interactions within the tumor microenvironment. Staining for mouse myeloid lineage cells (CD45, F4-80, CD11b, CD11c, GR-1) detected a minor population of mouse macrophages in the xenograft tumor (average ∼3% of viable cells (not shown), compared to the expected proportion of ∼30-50% macrophages in human GBM^26^). Thus, the immune suppressive macrophage population that is abundant in human GBM may not play as significant a role in modulating the effectiveness of CAR-T cell therapy in this model.

Next, tissue-infiltrating CAR-T cells were directly assessed by flow cytometry using the anti-idiotypic 1A7 antibody specific for the CAR antigen binding domain^33^. CAR-T cells were detected in brain, spleen, and bone marrow by flow cytometry, confirming successful homing and engraftment (**Figure 4D and Supplementary Figure 5B**). We noted that GBM patient-derived CAR-T cell therapy resulted in significantly higher absolute numbers of CAR-T cells in the brain, although this did not result in improved survival in these mice compared to mice treated with healthy donor-derived CAR-T cells.

To further evaluate the association of these findings with survival of tumor-bearing mice, treated mice were split into two groups based on survival times (short- and long-term survivors, see **Supplementary Figure 5A**). Absolute CAR-T cell numbers varied substantially within each group, but overall, higher numbers of CD3^+^, CD4^+^ and CD8^+^ CAR-T cells within the brain were significantly associated with longer survival **(Supplementary Figure 5C**). Absolute numbers of CD3^+^ CAR-T cells in spleen and bone marrow were less strongly linked to survival **(Supplementary Figure 5D-E**), however long-term survivors had significantly higher numbers of CD3^+^ CAR-T cells in bone marrow.

Thus, GD2-CAR-T cells were able to engraft peripherally and access the brain tumor, and these finding were linked with longer overall survival in tumor-bearing mice.

### Expression of an IL-15 transgene significantly improves CAR-T engraftment and tumor control

Given that CAR-T cell engraftment is a known and strong correlate of its clinical effectiveness^34^, and that we found CAR-T cell engraftment and infiltration of the brain correlated with improved survival, next we investigated ways to enhance these factors. A straightforward experimental approach would be to increase the CAR-T cell dose in mice (i.e. from the 10^6^ range to the 10^7^ range) but this would exceed the equivalent cell dose that would be practicable in patients^35^. Alternatively, administration of a similar dose of CAR-T cells engineered to have a greater proliferative potential may achieve the same end as a higher cell dose.

To investigate this possibility, we generated GD2-CAR-T cells using another retrovector, which incorporated a CAR transgene encoding a CD28-CD3ζ T-cell endodomain, ad additional transgene encoding secreted human IL-15. as well as other structural differences shown in **Supplementary Figure 6A** (kindly supplied via the Baylor college of Medicine, *in vitro* characterization shown in **Supplementary Figure 6**)^36,37,^. Like the third-generation GD2-CAR retrovector that we and our collaborators have used in the CARPETS and GRAIN clinical studies^16,17^, this second-generation GD2-CAR-IL-15 retrovector is being used in an active clinical study (clinicaltrials.gov: NCT03294954; NCT03721068).

When mice were treated with healthy donor-derived GD2-CAR-T cells enhanced with the IL-15 transgene, highly effective tumor control was observed, with a mean survival of 63.5 days (**Figure 5**). Mice treated with the third-generation GD2-CAR-T cells had a mean survival of 47 days and, similar to the previous experiment, the mean survival times for control mice were 41 and 42 days for untreated and NT-T cell treated mice, respectively (**Figure 5A-B**). In 3 of 6 mice treated with GD2-CAR-IL-15-T cells, the bioluminescence signal was undetectable at 49-63 days. However, at the final time point (day 67), a recrudescent low-level bioluminescence signal suggested tumor re-growth. Therefore, all remaining mice were culled to allow an *ex vivo* assessment of tumor burden and CAR-T cell infiltration. As GD2-CAR-IL-15-T cells were more often culled because of poor body condition rather than neurological signs (**Figure 5C**) and had markedly enlarged spleens (**Supplementary Figure 7A**), we suspected GvHD, as reported by others^18^. As with the previous experiment, CAR-T cell treated mice did not show early onset of neurological signs compared to control mice, and there was no evidence of off-tumor CAR-T cell-mediated neurotoxicity (**Figure 5C**). Brain tissues from two individual mice from each treatment group were used for bulk next generation RNA sequencing to provide a transcript-level overview of changes in the tumor microenvironment between groups (**Figure 5D**). GD2-CAR-IL-15-T cell-treated mice had higher levels of some chemokine receptors (CCR5, CX3CR1) and markers associated with cytotoxic function (PRF and GZMs) compared to GD2-CAR-T cell-treated mice, indicating possible differences in homing and cytolytic potential between these two clinically relevant GD2-CAR-T cell products. The mice treated with the third-generation GD2-CAR-T cells had notably higher levels of markers associated with a vasculature permissive to T-cell extravasation.

**Figure 5.**
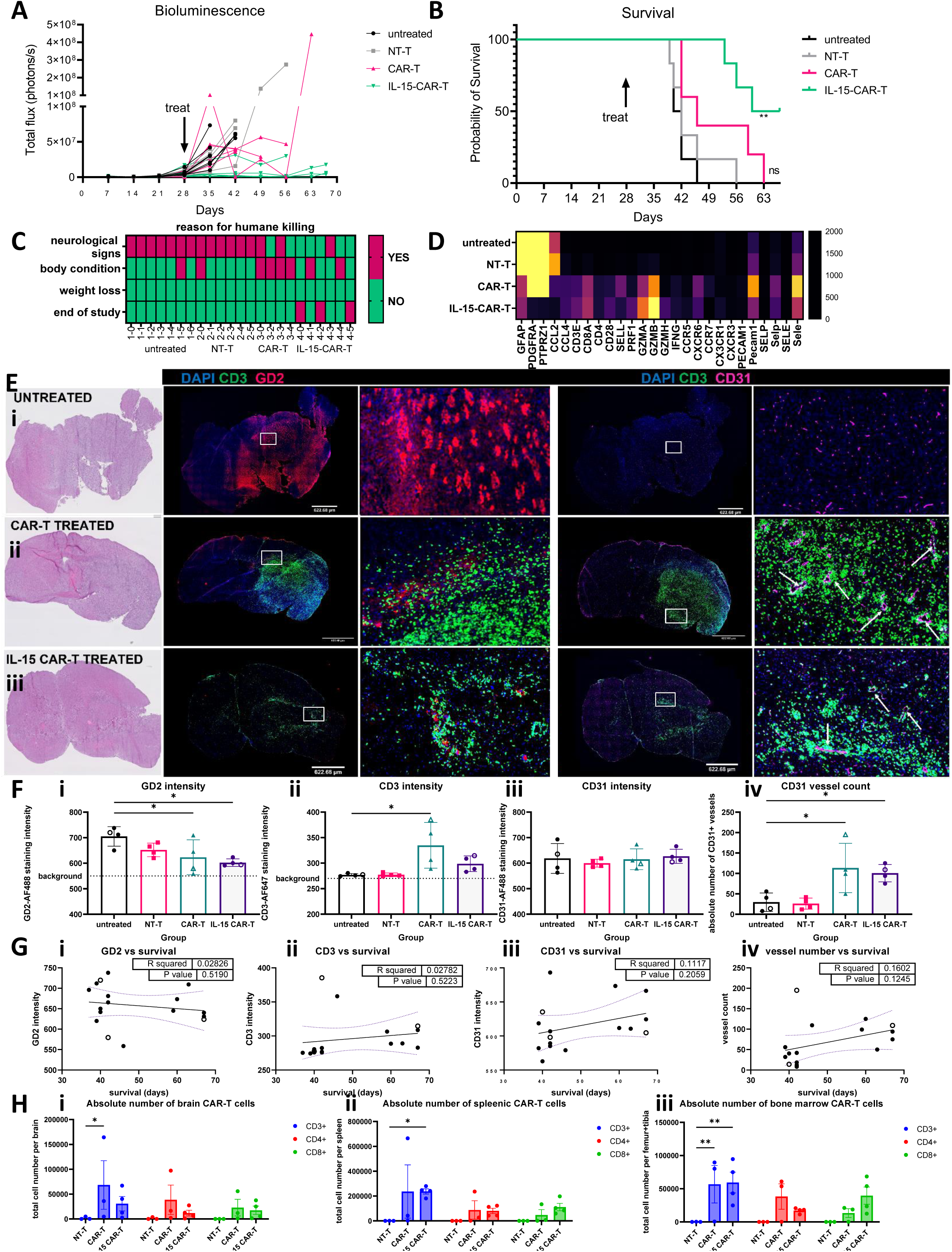
Incorporating an IL-15 transgene confers superior orthotopic tumor control by second-generation GD2-CAR-T cells compared to third-generation GD2-CAR-T cells. Mice received 2 × 10^5^ CCB-G5c cells by stereotactic injection on Day 1. On Day 28, mice were given single intravenous injections of saline (untreated) 1.5 × 10^6^ non-transduced control T cells (NT-T) from healthy donor, 3 × 10^6^ healthy donor-derived third-generation GD2-CAR T cells (CAR-T) or 3 × 10^6^ IL-15-containing GD2-specific CAR-T cells (IL-15-CAR-T). *n* = 5-6/group **A)** BLI data for all mice. **B)** Kaplan-Meir survival curves and statistics for mice. Statistics shown are from individual Gehan-Breslow-Wilcoxon tests comparing each curve to the untreated curve. **C)** Clinical signs resulting in humane killing according to pre-defined criteria (See methods). **D)** Next-generation sequencing of whole brain dissociated tissues from 2 untreated, 2 NT-T treated, 2 CAR-T treated and 2 IL-15-CAR-T treated mice. Heat map of selected markers show average normalized mRNA counts (Reads per KB per million; RPKM). **E)** Whole brain sections (mid-coronal plane where possible) were assessed by H&E staining (far left column), and immunofluorescence (IF) using anti-GD2-AF647 (clone 14g2a) and anti-human CD3 AF488 (columns 2 and 3) or anti-human CD3 AF488 and anti-mouse CD31-AF647 (columns 4 and 5) antibodies. White boxes on the whole brain sections indicate regions of interest shown at higher magnification to the right. White arrows indicate large vessels, identified by the presence of a black lumen and distinct from microvessels. Representative images from an i) untreated mouse, ii) CAR-T treated mouse, and iii) an IL-15-containing CAR-T treated mouse have been chosen to show the range of expression for each molecule. *n* = 4/group **F)** ImageJ analysis of the staining intensity for each antibody (i-iii), and enumeration of the number of large CD31^+^ vessels (iv). The individual mice represented in the IF microscopy images are shown with open symbols. **G)** Linear regression analysis of IF staining and survival. **H)** Enumeration of CAR^+^ T cells by flow cytometry from dissociated brain, spleen, and bone marrow, *n* = 4/group.

At study endpoints, GD2 staining was high in the tumors of control mice (untreated and NT-T treated), and lower in all CAR-T treated mice (**Figure 5E-F**), and the GD2-CAR-IL-15-T treated mice also had lower GFAP staining, consistent with the observed reduction in bioluminescence signals and indicating significantly decreased tumor volumes (**Figure 6**). The CD3 staining intensity in brain was highest for third-generation GD2-CAR-T treated mice. However, unlike in the previous experiment, a linear regression analysis showed that the CD3 staining intensity did not correlate with survival because in this experiment long-term survivors among GD2-CAR-IL-15-T treated mice had intermediate intensity of CD3 staining (**Figure 5F-G)**. While CD31^+^ vessels were again highest among of the CAR-T treated mice, mice with higher numbers of CD31 vessels did not always survive longest, unlike in the previous experiment (**Figure 5F-G**).

**Figure 6.**
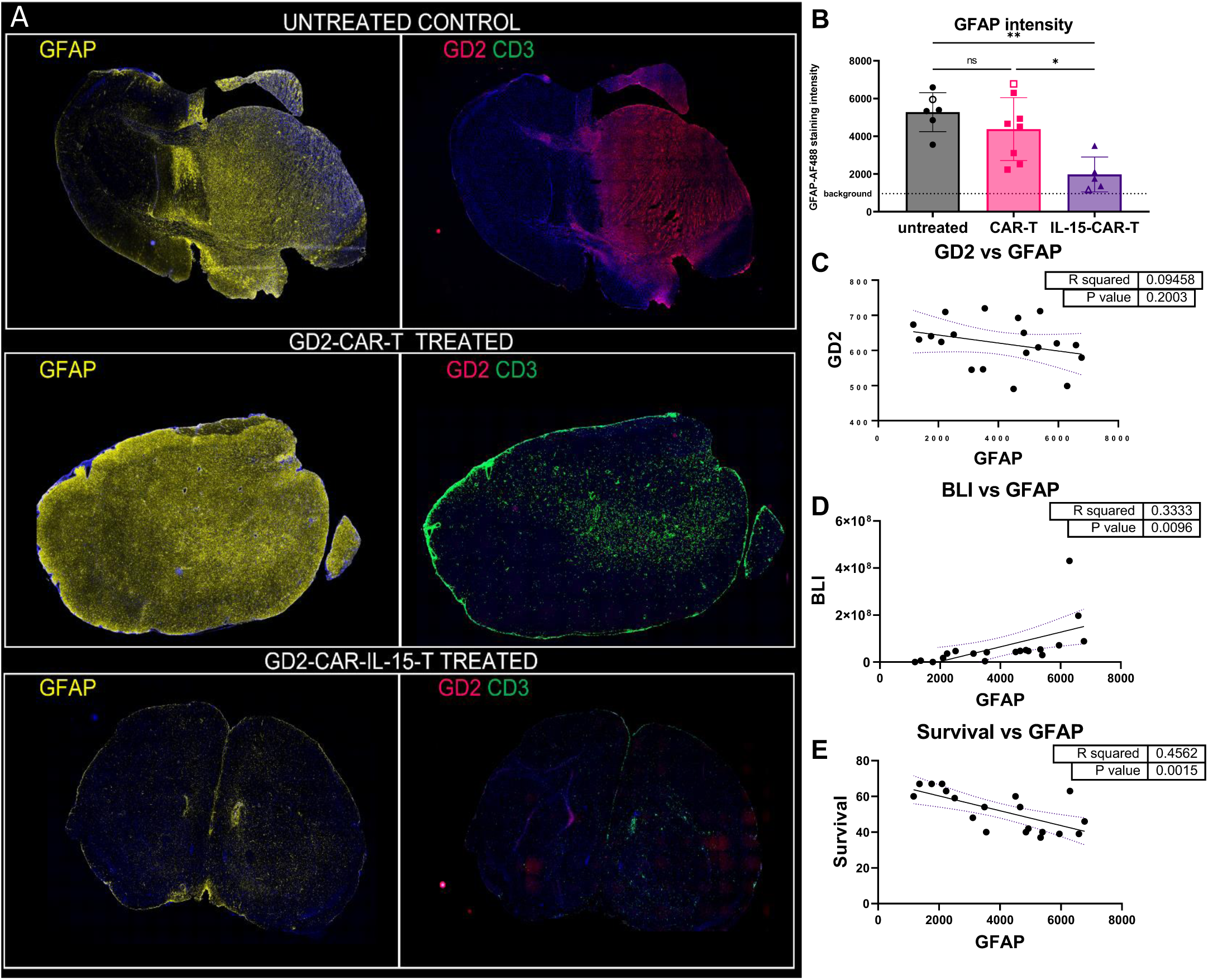
Staining for human GFAP reveals the extent of tumor control vs. tumor escape.. **A**) Whole brain sections were stained with rabbit anti-human GFAP. Contiguous sections stained with GD2 and CD3 are presented alongside to enable direct comparison. Representative images from i) untreated mouse, ii) GD2-CAR-T treated mouse, and iii) GD2-CAR-IL-15 treated mouse. *n* = 6-8/group. See also **Supplementary table 3** for a full data summary. **B)** ImageJ analysis of the staining intensity for human GFAP antibody. The individual mice represented in the IF microscopy images are shown with open symbols. Linear regression analysis of GFAP IF staining and **C)** GD2 staining intensity **D**) bioluminescence (BLI - total flux) at end-point **E)** survival (days).

Flow cytometry analysis of dissociated tissues confirmed the presence of GD2-CAR-expressing T cells in the brain, spleen, and bone marrow (**Figure 5H and Supplementary Figure 7C**). As we had observed in the CD3 immunofluorescence analysis, the third-generation GD2-CAR-T treated mice had significantly higher numbers of CAR-positive T cells in the brain compared to the GD2-CAR-IL-15-T treated mice. The latter mice had significantly higher numbers of CAR-positive T cells in the spleen, although the numbers of CAR-positive T cells in bone marrow in both groups of CAR-T treated mice were equivalent.

Together these data suggest that the two different GD2-CAR-T cell products have different engraftment and homing potentials. Compared to the third-generation GD2-CAR-T cells, the antitumor effectiveness GD2-CAR-IL-15-T cells appears to depend less on absolute numbers and suggests that these CAR-T cells have a distinct biology and employ unique mechanisms to achieve tumor control.

### Immune phenotypic changes in CAR-T cells following infusion

To better understand the biology of CAR-T cells after infusion, and to identify potential hurdles to achieving lasting tumor control, we performed a high-parameter flow cytometry analysis of single cell suspensions of spleen, bone marrow, and brain (**Figure 7** and gating strategy in **Supplementary Figure 8)**. This panel included T-cell subset markers (CD3, CD4, CD8, CD45RA, CD28), the anti-CAR 1A7 antibody^33^, markers of activation and exhaustion (PD-1 and LAG-3), and chemokine receptors (CCR7, CCR5, CXCR6, CX3CR1), which were selected because of their link to T-cell differentiation and glioblastoma homing^38^, and their differential expression in the RNAseq analysis described above.

**Figure 7.**
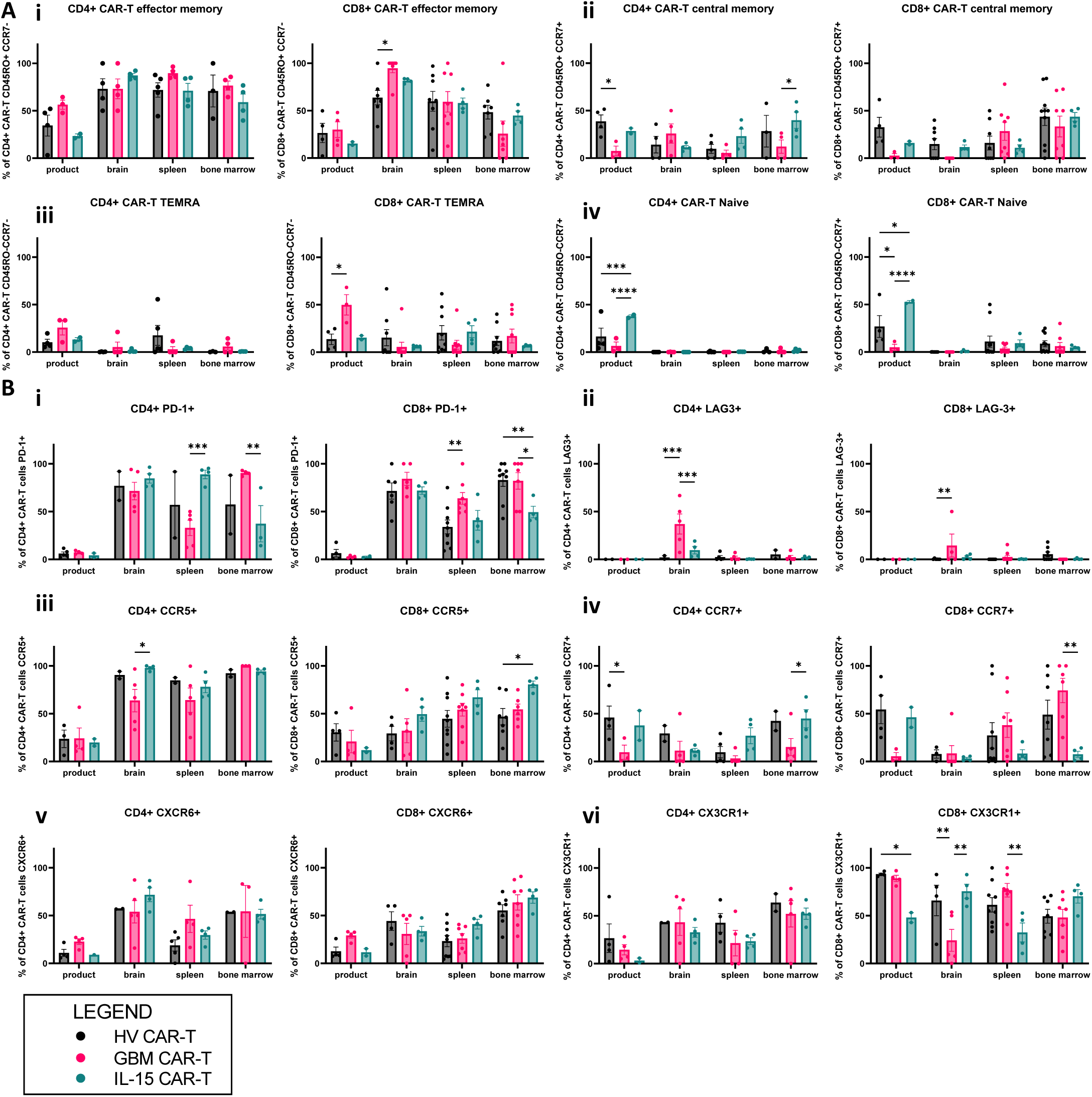
Multicolor flow cytometric analysis of brain-infiltrating, splenic and bone-marrow resident CAR T cells. A 14-color antibody panel (see **Supplementary Figure 9** for gating strategy and representative dot plots and **Supplementary Table 2** for antibody details) was designed to assess T-cell surface markers that might associate with CAR-T cell homing, persistence and efficacy and used to stain single cell suspensions of mouse brain, spleen, and bone marrow (right femur and tibia). Only mice with sufficient CAR-T cells were included in analysis, defined as at least 10^8^ live CAR^+^ CD3^+^ cells, which was the average of counts for control mice (background) + three standard deviations. Of the 30 CAR-T treated mice in three independent experiments, which included third-generation and IL-15-expressing second-generation GD2-CAR-T cells of healthy and GBM patient donor origin, 26 mice had tissues suitable for analysis by flow cytometry. Of these 26 mice, 11 and 21 mice had enough CAR^+^ CD4^+^ and CD8^+^ T cells, respectively, for the subset analyses. **A)** Memory phenotype on CAR-T cell products and CAR-T cells in the brain, spleen, and bone marrow i) CD45RO^+^ CCR7^-^ TEM, ii) CD45RO^+^ CCR7^+^ TCM, iii) CD45RO^-^ CCR7^-^ TEMRA, iv) CD45RO^-^ CCR7^+^ Naïve T cells. **B)** Expression of T-cell surface markers of interest on CAR-T cell products and CAR-T cells in the brain, spleen, and bone marrow. i) PD-1 on CD4^+^ (left) and CD8^+^ CAR-T cells, ii) LAG-3 on CD4^+^ (left) and CD8^+^ CAR-T cells, iii) CCR5 on CD4^+^ (left) and CD8^+^ CAR-T cells, iv) CCR7 on CD4^+^ (left) and CD8^+^ CAR-T cells, v) CXCR6 on CD4^+^ (left) and CD8^+^ CAR-T cells, vi) CX3CR1 on CD4^+^ (left) and CD8^+^ CAR-T cells.

Expression of the markers of interest was broadly similar among the starting products (**Figure 7**). GBM patient-derived CAR-T cells had less CCR7 expression and a less naïve or central memory-like phenotype, and more TEMRA-phenotype T cells compared to healthy donor products. The GD2-CAR-IL-15-T cells contained a higher proportion with a naïve or central memory phenotype, as reported previously^39^. After infusion, the dominant CAR-T cell population observed in mice of all groups was of an effector memory phenotype in the brain, spleen, and bone marrow (**Figure 7A**). GD2-CAR-IL-15-T cells contained a higher proportion of central memory CD4^+^ CAR-T cells in the bone marrow.

All CAR-T cells upregulated PD-1 compared to the starting product, which had very low PD-1 expression, indicating activation of these cells (**Figure 7B i**). In contrast, LAG-3 was negative on the CAR-T cell products and not significantly upregulated on CAR-T cells in the spleen or bone marrow. LAG-3 was significantly upregulated on the brain CD4^+^ CAR-T cells, but only for CAR-T cells from GBM donors, indicating that products from these patients may be inherently more exhausted or susceptible to suppression (**Figure 7B ii)**.

Expression of chemokine receptor CCR5, which is a marker of differentiation from naïve to effector memory and which contributes to T-cell homing to tumors^40^, was significantly upregulated on all tissue-homing CAR-T cells and, compared to the third-generation GD2-CAR-T cells, GD2-CAR-IL-15-T cells had the highest proportion of CCR5^+^ cells in the brain and bone marrow (**Figure 7B iii**). In most cases, the proportion of cells expressing CCR7 was not significantly different to the infusion product. However, in the bone marrow, CD4^+^ GD2-CAR-IL-15-T cells had significantly higher CCR7 expression and CD8^+^ GD2-IL-15-CAR-T cells had significantly lower CCR7 expression than the equivalent GBM-derived CAR-T cells, which have the opposite phenotype (**Figure 7B iv**). CXCR6, which is involved in localizing T cells to the tumor^41^, was upregulated in all tissue CAR-T cells compared to pre-infusion CAR-T cell products, with no differences observed among treatment groups (**Figure 7B v**). CX3CR1 is considered a marker of differentiation and although CX3CR1-negative T cells are generally less differentiated and more polyfunctional in the periphery, T cells lacking CX3CR1 in the tumor have poorer function and are more susceptible to immune suppression via checkpoint molecules^42^. In the pre-infusion product, the GD2-CAR-IL-15-T cells had lower expression of CX3CR1, consistent with a less differentiated phenotype. In tissue, however, there was a significantly lower proportion of CX3CR1^+^ CD8^+^ GD2-CAR-T cells in the brain of mice treated with GBM-patient derived CAR-T cells compared to healthy donor-derived CAR-T cells (**Figure 7B vi**).

Thus CCR5, CXCR6 and CX3CR1 were all strongly expressed on tissue-localized CAR-T T cells, and GBM-derived CAR-T cells had higerh levels of exhaustion markers. Although interesting differences in CAR-T cell phenotype among individual mice were observed, we could not determine if there was a correlation between specific CAR-T cell phenotypes in tissues and overall survival as long-term survivors were more likely to have enough CAR-T cells for further analyses compared to short-term survivors, which are thus not sufficiently represented in the phenotypic analyses.

## Discussion

This is the first study to validate a clinically relevant GD2-CAR-T cell product as a therapy for GBM. Importantly, in this study we have used patients’ blood samples for CAR-T cell manufacture and patients’ tissues to establish early-passage glioma neural stem cell lines as CAR-T cell targets and to generate orthotopic tumors. Hence, in contrast to prior preclinical studies using tissue culture-established tumor cell lines and healthy donor-derived CAR-T cells, we have evaluated this CAR-T cell therapy by also considering the state of the tumor and immune compartments in GBM patients.

GD2 has long been a tumor-associated antigen of interest^12^. Here, we provide a comprehensive survey of GD2 expression using primary tissue samples obtained from GBM and DMG patients, revealing high-level GD2 expression on surgical biopsies. However, as with all tumor-associated antigens, care must always be taken to consider patterns of expression on healthy tissues because of the known on-target, off-tumor toxicities of CAR-T cell therapy. As a normal tissue antigen, the neuronal expression of GD2 is an apparent concern for the treatment of GD2^+^ malignancies in the brain, which is encased in the confined space of the skull and thus less able to tolerate potential immuno-inflammatory reactions of CAR-T cell therapy.

As the donor of the same antigen binding region as US FDA-approved dinutuximab^15^, the 14g2a mAb also supplies the scFvs employed in GD2-CARs of this study. Of note, the GD2 epitope of the 14g2a-derived scFv is identical between human and mouse. The 14g2a-derived scFv has low affinity binding for GD2 in contrast to another GD2-specific and clinically studied mAb, 3F8^43^. Preclinical murine studies illustrate the clinical significance of GD2-specific scFv affinity^32^ as on-target, off-tumor lethal neurotoxicity associated with CAR-T cell infiltration and neuronal destruction was observed in mice administered GD2-CAR-T cells incorporating either a higher affinity mutation in the antigen-binding region of the 14g2a-derived scFv^14^ or a 3F8-derived scFv. Importantly, however, such toxicity was not observed in mice administered GD2-CAR-T cells containing the unmodified 14g2a-derived scFv^32^. In a preclinical murine study of DMG in which mice were administered GD2-CAR-T cells containing the unmodified 14g2a-derived scFv, on-target, on-tumor neurotoxicity was observed in the absence of on-target, off-tumor neurotoxicity^18^. This neurotoxicity was attributed to tumor-related inflammation and edema without evidence of brain parenchymal CAR-T cell infiltration and neuronal destruction and proved lethal because of space-occupying effects when the tumors were in the thalamus^18^.

We have shown levels of GD2 expression in glioblastoma tissues that clearly exceed that detected in normal brain confirming the preclinical findings above^32,18^, and an independent pathologist reported no evidence of off-tumor GD2-CAR-T cell toxicity in the orthotopic xenograft GBM model. Indeed, GD2-CAR-T treated mice survived significantly longer than their control counterparts with euthanasia endpoints of poor body condition or excessive weight-loss consistent with the GvHD reported by others^18^ were reached later than the neurological endpoint reached more often in control mice (**Figure 3C-D and 5B-C**).

The observed lag in the intracranial bioluminescence signal in mice treated with third-generation GD2-CAR-T cells indicated transient tumor control before eventual tumor escape with onset of clinical signs. However, using another clinically studied GD2-CAR retrovector encoding the IL-15 transgene, we generated CAR-T cells that resulted in more complete and sustained tumor control. The GD2-CAR-IL-15-T cells further prolonged survival with a 50% complete response rate at 4-weeks post treatment as assessed by bioluminescence intensity. All mice from this group had GD2^lo/negative^ residual tumors at necropsy whereas 60% of GD2-CAR T cell treated mice still had substantial GD2^+^ tumors (**Figure 5A** and **Supplementary Table 3**). Importantly the GD2-CAR-IL-15 treated group also had significantly lower human GFAP staining in the brain, indicating there was not significant regrowth of GD2-negative GFAP+ tumor (**Figure 6**).

Our results differ from a recent study employing an orthotopic GBM model of xenografts and a second-generation CAR construct based on a novel IgM-derived and human GD2-specific scFv lacking cross-reactivity with murine GD2. The resulting GD2-CAR-T cells had significant antitumor activity only after intracerebral administration^23^. In contrast, in our study, the GD2-CAR-T cells were active against orthotopic glioblastoma xenografts after intravenous administration, perhaps because of the design of our CAR^44^ or of our optimized manufacturing process^27^.

Immunofluorescence staining and analysis of mouse brains at study endpoints suggested that tumors escaped CAR-T control either because of unrelenting immune selection pressure resulting in antigen loss (residual GD2^−^, hGFAP^+^ tumors) or intrinsic CAR-T cell defects (residual GD2^+^ tumors) (**Figure 6** and **Supplementary Table 3**). We hypothesize that the third-generation CAR-T cell therapy could not kill all antigen-positive tumor cells because of an inadequate initial cell dose, poor tumor infiltration by T cells, or inadequate T-cell functions such as cytotoxicity at the tumor site. The longest surviving mice had the highest levels of CD3^+^ T cells, as measured by CD3 tissue staining intensity and absolute numbers as counted by flow cytometry, and the greatest numbers of large intratumoral CD31^+^ blood vessels. Total CD31 staining intensity however did not significantly correlate with survival, suggesting that the CD31^+^ tumor microvessels might not support CD3 T-cell infiltration. This finding raises the possibility that tumor vasculature-normalizing effects of bevacizumab co-treatment may promote T-cell infiltration^45^.

IL-15 co-expression significantly increased GD2-CAR-T cell engraftment in the spleen and bone marrow of tumor-bearing mice although the improved survival was not simply related to a higher CAR-T cell dose reaching the tumor. Indeed, GD2-CAR-IL-15-T cells were equivalent or even less abundant in the brain compared to third-generation GD2-CAR-T cells but nonetheless exerted superior tumor control, as measured by all 3 of our tumor markers: GD2, bioluminescence and human GFAP. We hypothesize that the increased effectiveness of the GD2-CAR-IL-15-T cells reflected greater cytotoxic potency or improved survival in the GBM microenvironment or both. Others have reported a distinct phenotype that allows these IL-15-expressing-CAR-T cells to better survive repeated tumor challenge in a neuroblastoma model^36^. Our own RNA sequencing identified multiple differentially expressed pathways associated with chemokine responses and cytotoxicity that were elevated for GD2-CAR-IL-15-T treated mice.

High-parameter flow cytometry revealed that mice with the best post-treatment survival had significantly higher numbers of CD4^+^ and CD8^+^ CAR-T cells in the brain, and total CD3^+^ CAR-T cells in bone marrow. We were not able to uncover the CAR-T cell phenotype that associated with superior engraftment or infiltration, although the GD2-IL-15-CAR-T cells with superior tumor control did have some distinct differences in phenotype including a higher proportion of bone-marrow CD4+ central memory CAR-T cells with lower relative PD-1 expression, and in the brain more CD4+ CAR-T cells expressing CCR5 and more CD8+ CAR-T cells expressing CX3CR1. This remains an area for further investigation. For all CAR-T treated mice, the receptors CCR5, CXCR6, CX3CR1 were highly represented on brain-infiltrating CAR-T cells, highlighting a role in T-cell recruitment described in other models^40-42^, and in our recent analysis of endogenous T cells in GBM^38^. We also noted that tumor-infiltrating CAR T cells were almost uniformly PD-1^+^, but GBM-patient derived CAR-T had elevated LAG-3 and were more likely to be CX3CR1-negative, indicating that they may be more susceptible to immune-suppression^42^.

Other factors that may impair the effectiveness of CAR-T cell therapy include known tumor microenvironmental inhibitors of T-cell function. The GNS cell line employed in the model showed a range of immune-modulatory molecules (IL-6, PD-L1, **Supplementary Figure 9**) which may interfere with T-cell function^46^. However, in human GBM xenografts hosted in the brains of NSG mice, and unlike in humans^38^, lymphocytes are absent and microglia and glioma-associated macrophages (GAMs) are relatively lacking, thus indicating that this orthotopic xenograft model does not adequately recapitulate the complex GBM microenvironment in patients.

As well as increasing our understanding of GD2-CAR-T cell behaviour in orthotopic xenografts of glioblastoma, we have been able to assess the practical aspects of manufacturing CAR-T products for glioblastoma patients. We identified technical hurdles to autologous CAR-T cell therapy for glioblastoma patients including a low yield of lymphocytes together with a skewed CD4:CD8 lymphocyte ratio and lower proportions of naïve T cells, which have been reported previously^31^. Nevertheless, potent *in vitro* functions were observed, and *in vivo* there was no significant difference in survival between mice treated with healthy-donor derived CAR-T cells or GBM patient-derived CAR-T cells.

Our advanced planning for a phase 1 trial of GD2-CAR-T cell therapy in glioma patients has caused us to consider the timing of CAR-T cell manufacturing and administration with respect to standard treatments. For example, we propose to obtain apheresis products from DMG patients before commencement of palliative radiotherapy, which aims to delay disease progression until the CAR-T cell product is ready for administration. Moreover, given the key role of blood vessels in T-cell infiltration of tumors we also propose to use the anti-VEGF-neutralizing mAb, bevacizumab, in combination with GD2-CAR-T cell therapy to mitigate brain swelling from CAR-T cell related neuroinflammation and potentially to stabilise the tumor vasculature so that T-cell access to the tumor is promoted^47-49^.

In summary, by using primary patient samples for manufacturing and disease modeling in this preclinical study, we have identified several factors associated with the duration of tumor control and ultimately the survival of glioblastoma-bearing mice. Successful CAR-T cell engraftment in bone marrow, high absolute numbers of tumor-infiltrating CAR-T cells and the development of a T cell-supportive intratumoral vasculature were all associated with improved survival following treatment with third-generation GD2-CAR-T cells that will be used in our upcoming phase 1 trials. Treatment with GD2-CAR-T cells enhanced with an IL-15 transgene had improved engraftment and significantly better tumor control, and without an evident increase in the number of tumor-infiltrating CAR-T cells. Our results support the case for clinical investigation of this approach albeit with the recognition that additional safety precautions such as incorporation of a suicide gene may be required were an IL-15 transgene able to promote autonomous CAR-T cell proliferation^37^.

## Materials and Methods

Detailed methodology provided in **Supplementary Methods**

### Human tumor material

Glioblastoma patient tumor tissue and blood were obtained through the South Australian Neurological Tumor Bank (SANTB). Use was approved by the Central Adelaide Local Health Network Human Research Ethics Committee (CALHN HREC; approval number R20160727). Melanoma patient blood samples for CAR-T cell manufacturing were obtained under the CARPETS trial protocol (CALHN HREC; approval number R20100524). Sample from DIPG patients were obtained from the Zero Childhood Cancer Initiative, and from the Sydney Children’s Hospital (SCHN HREC; 2019/ETH05438). Fresh tissue samples were prepared as described previously^26^. Peripheral T cells were separated from blood samples and used to generate GD2-specific third-generation CAR-T cells using GD2-iCAR retroviral vector SFG.iCasp9.2A.14g2a.CD28.OX40.zeta supernatant (Center for Cell and Gene Therapy, Baylor College of Medicine, Houston, TX, USA) and our clinical manufacturing protocol as described previously^27^. The second-generation GD2-CAR-IL-15 retrovector^36,50,51^ was obtained under a material transfer agreement with Baylor College of Medicine with the kind assistance of Prof. Leonid Metelitsa. All human specimens were used in accordance with the Declaration of Helsinki, and participants provided written consent.

### Culture of GNS tumor cell lines and CAR-T cells

Glioma neural stem (GNS) cells were cultured in StemPro NSC medium (Thermo Fisher) as described previously^26^. The neuroblastoma line LAN-1 was maintained in DMEM-F12 media with 10% FCS, 1% Glutamax, 1% Pen/Step. Post-thawing, CAR-T cell products were maintained in TEXMACS media (Miltenyi) with 10ng/mL IL-7 and 5ng/mL IL-15 for a maximum of 10 days.

### Immunofluorescence staining of cryosections

Sections were cut from OCT-embedded human glioblastoma, human adjacent normal brain, and mouse brain tissues (5-6µm thickness).

Tissue sections were stained using 5µg/mL purified mouse anti-CD3 (UCHT1; BioLegend) and 4µg/mL purified hamster anti-CD31 (2H8; Thermo Fisher Scientific) with goat anti-mouse IgG AF647 (Thermo Fisher Scientific) and goat anti-hamster IgG AF488 (Abcam) detection antibodies, and mouse anti-GD2-FITC (14.G2A; BD) at a dilution of 1:100. GFAP was detected with polyclonal rabbit-anti human GFAP (1:500, Agilent) and donkey anti-rabbit IgG AF55 detection (1:500, Thermo Fisher Scientific).

Whole-slide imaging was performed on a Zeiss Axio Scan.Z1 slide-scanner using 20x objective and ZEN 3.1 Blue system software. Fluorescence overlays were created by merging channels and applying false color using FIJI (ImageJ, National Institutes of Health).

For further detail see **Supplementary methods**.

### Flow Cytometry

Stained cells were analyzed on a BD LSR Fortessa (BD Biosciences) or a BD FACSymphony A5 (BD Biosciences) with FlowJo Software V 10.8.1 (BD Biosciences). For further detail see **Supplementary methods**. Antibodies for multicolor panels are listed in **Supplementary Table 2**.

### Orthotopic Xenograft Model

Animal experiments were conducted under a protocol approved by the University of South Australia Animal Ethics Committee (#U44-19). A Stoelting motorized stereotactic alignment and injection unit was used to deliver 2μL of patient-derived GNS cells (2-5×10^5^ cells) over 4 minutes, 3mm deep into the right hemisphere of 6-8 week-old female NOD-SCID-gamma-null (NSG) mice. Mice received daily clinical checks and weekly bioluminescence imaging to monitor tumor growth. Bioluminescence imaging was performed using an IVIS Lumina S5 after intraperitoneal injection of 100μL of 30mg/mL luciferin in saline.

CAR-T cell treatment (up to 1×10^7^ total T cells; 1.5-4 ×10^6^ CAR^+^ T cells) was administered intravenously *via* tail-vein injection once tumor growth had been observed for two consecutive weeks, confirming tumor engraftment at days 17-28 in these experiments. Defined humane endpoints for euthanasia included loss of >15% body weight from starting weight, a body condition score of less than 2 (under-conditioned^52^), and neurological signs including head-tilt, loss of balance and circling movement.

### Statistics

Data were analyzed using GraphPad Prism Version 9.3.1. Single-variable data were analyzed by one-way ANOVA, two-variable date were analyzed by two-way ANOVA and Tukey’s multiple comparison post-tests. Multiple survival curves were analyzed using the logrank test for trend, and specific comparisons of two selected curves employed the Gehan-Breslow-Wilcoxon test. Statistical significance is represented on graphs as * ≤ 0.05, ** ≤ 0.01 and *** ≤ 0.001.

## Acknowledgments

We acknowledge Prof. Malcolm Brenner, Prof. Gianpietro Dotti, and A/Prof. Eric Yvon for provision of the original third-generation GD2-specific CAR-T cell retroviral vector. We acknowledge Prof. Leonid Metelitsa and Prof. Gianpietro Dotti for provision of the second-generation GD2-specific and IL-15 producing CAR retroviral vector. We acknowledge Prof. Bryan Day for GNS lines. We acknowledge Bryan Gardam for analysis of macrophage and microglia in NSG mice. We acknowledge Steven Roberts and Dr Andrew Lim (BD Biosciences) for technical advice in designing multi-color flow panels for use on the BD FACS Symphony. We acknowledge Dr Agatha Labrinidis for technical microscopy support. We acknowledge Miltenyi Biotec for support in sourcing the Clinimacs Plus instrument used in our trial manufacturing, and Paula Stoddart for technical advice. We acknowledge the Zero Childhood Cancer Initiative for access to archival DIPG patient material, and the KOALA clinical trials team (Emma McCormack, Lily Wong, Alison Rego) for recruitment of DIPG patients to give fresh blood samples. We also acknowledge the support and generosity of the patients and medical and technical staff from SA Pathology and the SA Neurological Tumor Bank (supported by Flinders University, Flinders Foundation and The NeuroSurgical Research Foundation), who made collection of tissue specimens possible. We thank clinical pathologists at SA Pathology and veterinary pathologists at Gribbles pathology (Adelaide) for their histopathologic evaluations. RNA-seq experiments were performed at the Australian Cancer Research Foundation (ACRF) Cancer Genomics and Cancer Discovery Accelerator facilities at SA Pathology, established with the generous support of the Australian Cancer Research Foundation, and bioinformatics support was kindly provided by John Toubia.

## List of Abbreviations

If abbreviations are used in the text, they should be defined in the text at first use, and a list of abbreviations should be provided.

CAR: chimeric antigen receptor
CCB: Centre for Cancer Biology, South Australia
DIPG: diffuse intrinsic pontine glioma
DMG: diffuse midline gliomas
GBM: glioblastoma
GD2: disialoganglioside; C_74_H_134_N_4_O_32_
GD2-CAR: third-generation GD2-specific CAR with inducible caspase-9 suicide gene
GD2-CAR-T: GD2-specific chimeric antigen receptor-bearing peripheral blood T cells
GD2-CAR-IL-15-T: Second-generation GD2-specific CAR with a transgene for secreted IL-15
GNS: glioma neural stem
mAb: monoclonal antibody
NSG: mouse strain NOD.Cg-Prkdcscid Il2rgtm1Wjl/SzJ
NT: matched, non-transduced patient T-cell control
PBMCs: peripheral blood mononuclear cells
PBT: peripheral blood T cells
retroviral vector: retrovector
scFv: single chain variable fragment
TCM: central memory T cells
TEM: effector memory T cells
TEMRA: effector memory T cells expressing CD45RA
TSCM: stem/central memory-like T cells

## Notes

**Competing Interests** The authors declare that the research was conducted in the absence of any commercial or financial relationships that could be construed as a potential conflict of interest.

**Funding** This work was supported by the NeuroSurgical Research Foundation, The Hospital Research Foundation Group, the Cancer Council SA Beat Cancer Project (Hospital Research Package and Fellowship to TG), the Health Services Charitable Gifts Board (Adelaide), Tour de Cure, the Ray & Shirl Norman Cancer Research Trust, the Mark Hughes Foundation, the National Health and Medical Research Council (Fellowship to SMP), and the Fay Fuller Foundation (Fellowship to MNT).

### Competing Interest Statement

The authors have declared no competing interest.

