## Supplementary Tables and Figures for "GD2-targeting CAR-T cells enhanced by transgenic IL-15 expression are an effective and clinically feasible therapy for glioblastoma"

### Supplementary Table 1. Donor characteristics and sample usage.

| Patient identifier | Brain tumour type | Treatment before blood donation | Steroids at blood donation | Demographics – sex, age (years) | GNS tumour cell line established? | CAR-T cell product manufactured? |
|---|---|---|---|---|---|---|
| **DIPG 1** | DIPG | Radiotherapy; chemotherapy ACT001 | None noted | Female; 3 | NA | yes |
| **DIPG 2** | DIPG | Radiotherapy | None noted | Female; 3 | NA | yes |
| **DIPG 3** | DIPG | Radiotherapy | None noted | Female; 4 | NA | yes |
| **DIPG 4** | DIPG | Radiotherapy | None noted | Male; 4 | NA | yes |
| **DIPG 5** | DIPG | Radiotherapy; cabozantinib | Yes | Male; 4 | NA | yes |
| **BT 11** | Recurrent GBM | Surgery; radiotherapy; TMZ | Yes | Not provided | Not established | yes |
| **BT 23** | Primary GBM | None | Yes | Male; 78 | CCB-G4 | no |
| **BT 26** | Primary GBM | None | Yes | Male; 69 | CCB-G5 | no |
| **BT 29** | Primary GBM; IDH WT | None | Yes | Male; 66 | CCB-G6 | yes |
| **BT 31** | Primary GBM; IDH WT | None | Yes | Male; 57 | CCB-G7 | yes |
| **BT 36** | Primary GBM | None | Yes | Male; 40 | CCB-G10 | yes |
| **BT 48** | Primary GBM | None | Yes | Male, 68 | Not established | yes |
| **BT B1** | Primary GBM | None | Yes | Male; 61 | Not established | yes |
| **CARPETS 101** | MM | None | No | Female; 54 | NA | yes |
| **CARPETS 102** | MM | None | No | Male; 72 | NA | yes |
| **CARPETS 201** | MM | None | No | Male; 62 | NA | yes |
| **CARPETS 203** | MM | None | No | Male; 62 | NA | yes |
| **CARPETS 304** | MM *with brain metastases | None | No | Male; 40 | NA | yes |
| **CARPETS 303 pre** | MM | None | No | Male; 45 | NA | yes |
| **CARPETS 303 post** | MM *with brain metastases | dabrafenib+ trametinib; ipilimumab +nivolumab | No | Male; 49 | NA | yes |

Abbreviations: diffuse intrinsic pontine glioma, DIPG; glioblastoma multiforme, GBM; metastatic melanoma, MM; not available, NA; temozolomide, TMZ

**Supplementary Table 2. Antibody Panels.**

| Basic immune phenotyping panel | 13 colour immune phenotyping panel |
|---|---|
| CD3 buv395 BD # | CD3 buv805 BD # |
| CD4 bv510 BD # | CD4 buv496 BD # |
| CD8 PeCy7 BD # | CD8 APCH7 BD # |
| CD45RA AF647 BD # | CD45RO bv480 BD # |
| CCR7 bv421 BD # | PD-1 PeCy7 #561272 |
| CD62L AF488 BD # | LAG-3 buv395 BD # |
| CD57 PE BD # | CD28 buv615 BD # |
| Fixable viability dye AF700 BD # | CCR7 bv786 BD # |
| | CCR5 buv737 BD # |
| | CXCR6 bv421 BD # |
| | CX3CR1 BB700 BD # |
| | L/D FVS575V BD # |
| | 1A7 anti-CAR antibody (in house) AF647 |

**Supplementary Table 3. Tumor characteristics at humane endpoint and necropsy.**

| Mouse | Treatment | CD3 staining | TUMOR MARKERS | | BLI signal | Survival (days) |
|---|---|---|---|---|---|---|
| | | | GD2 staining | hGFAP staining | | |
| 1 | untreated | none | high | high | high | 37 |
| 2 | untreated | none | low | low | high | 38 |
| 3 | untreated | none | low | low | high | 35 |
| 4 | untreated | none | high | ND | high | 39 |
| 5 | untreated | none | high | high | high | 40 |
| 6 | untreated | none | high | high | high | 40 |
| 1 | NT-T cell | none | high | high | high | 42 |
| 2 | NT-T cell | low | high | high | high | 39 |
| 3 | NT-T cell | none | high | high | high | 42 |
| 4 | NT-T cell | none | high | high | high | 40 |
| 1 | GD2-CAR T cell | low | high | high | high | 54 |
| 2 | GD2-CAR T cell | high | low | high | high | 60 |
| 3 | GD2-CAR T cell | high | low | high | high | 63 |
| 4 | GD2-CAR T cell | low | high | ND | high | 48 |
| 5 | GD2-CAR T cell | high | low | high | high | 42 |
| 6 | GD2-CAR T cell | high | low | high | high | 46 |
| 7 | GD2-CAR T cell | low | high | low | high | 63 |
| 8 | GD2-CAR T cell | high | low | high | high | 59 |
| 1 | GD2-CAR-IL-15 T cell | high | low | low | low | 67 |
| 2 | GD2-CAR-IL-15 T cell | high | low | low | low | 60 |
| 3 | GD2-CAR-IL-15 T cell | high | low | low | low | 67 |
| 4 | GD2-CAR-IL-15 T cell | high | low | low | low | 53 |
| 5 | GD2-CAR-IL-15 T cell | high | low | high | high | 54 |
| 6 | GD2-CAR-IL-15 T cell | high | low | ND | low | 67 |

**TABLE NOTES.** Summary data from 3 independent experiments, displayed in main figures 3 and 5. Only mice with high-quality fresh-frozen tissue samples were included in the staining analysis. Definitions: NT-T, non-transduced control T cells; ND, not determined. Staining intensity score was based on imageJ intensity measurements, with a high reading (red shading) defined as greater than 3x the standard deviation of the background signal (see main figures 4 and 5 for intensity graphs), and a low reading (blue shading) as anything below this value. Bioluminescence imaging (BLI) intensity scored low (blue shading) if total flux was below $1.9 \times 10^7$ (average signal from untreated control at endpoint – 2x standard deviations), with a high reading (red shading) as value above this level.

### Supplementary Figures

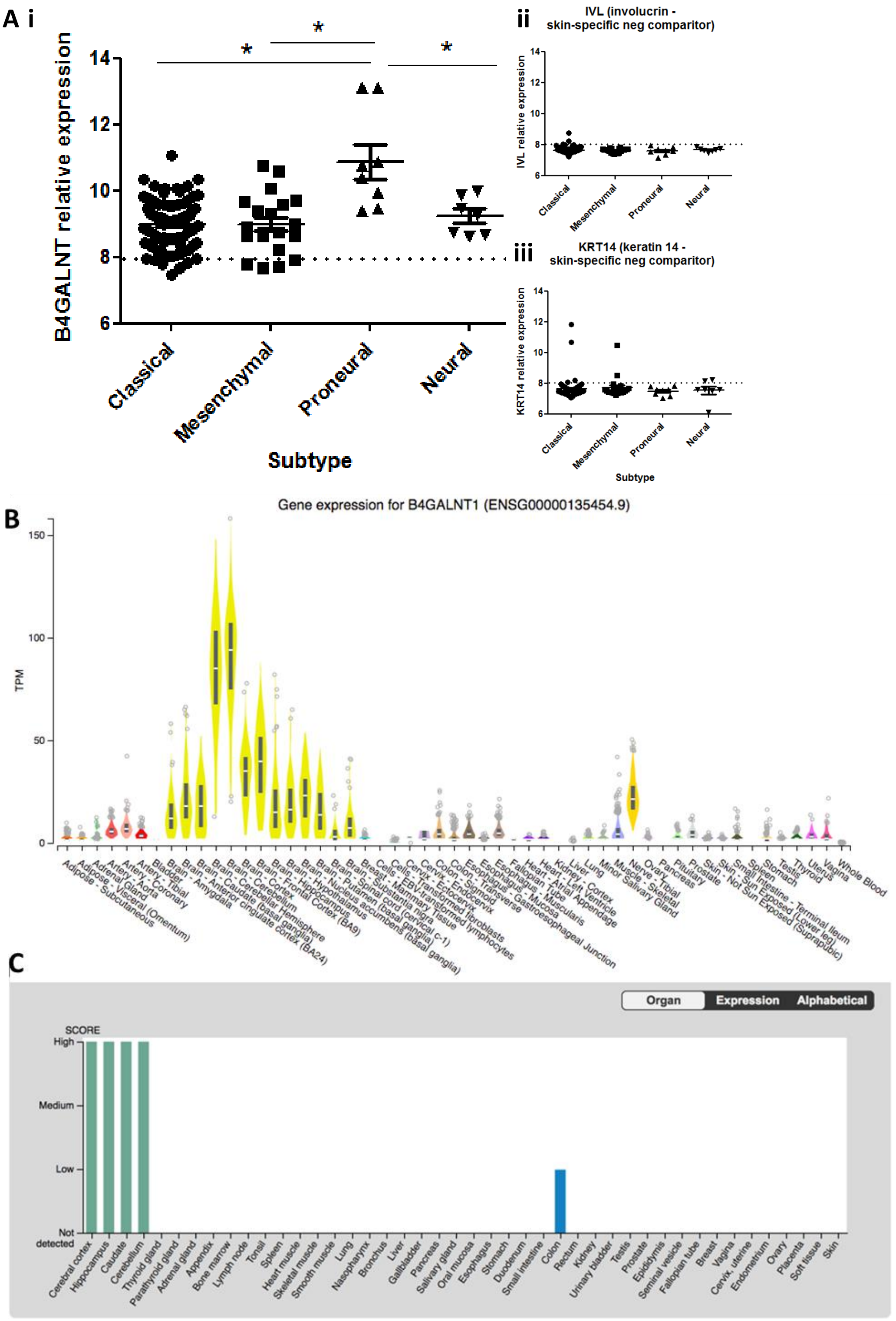


**Supplementary Figure 1. GD2 expression patterns in tumor and normal brain. A)** RNA expression of the GD2 synthase enzyme from Dataset GSE72951 - *Identification of recurrent GBM patients that may benefit from Bevacizumab and CCNU: A report from the BELOB trial by the Dutch Neurooncology Group.* **B)** RNA expression of the GD2 synthase enzyme. **C)** Protein expression of the GD2 synthase enzyme from the GTeX dataset.

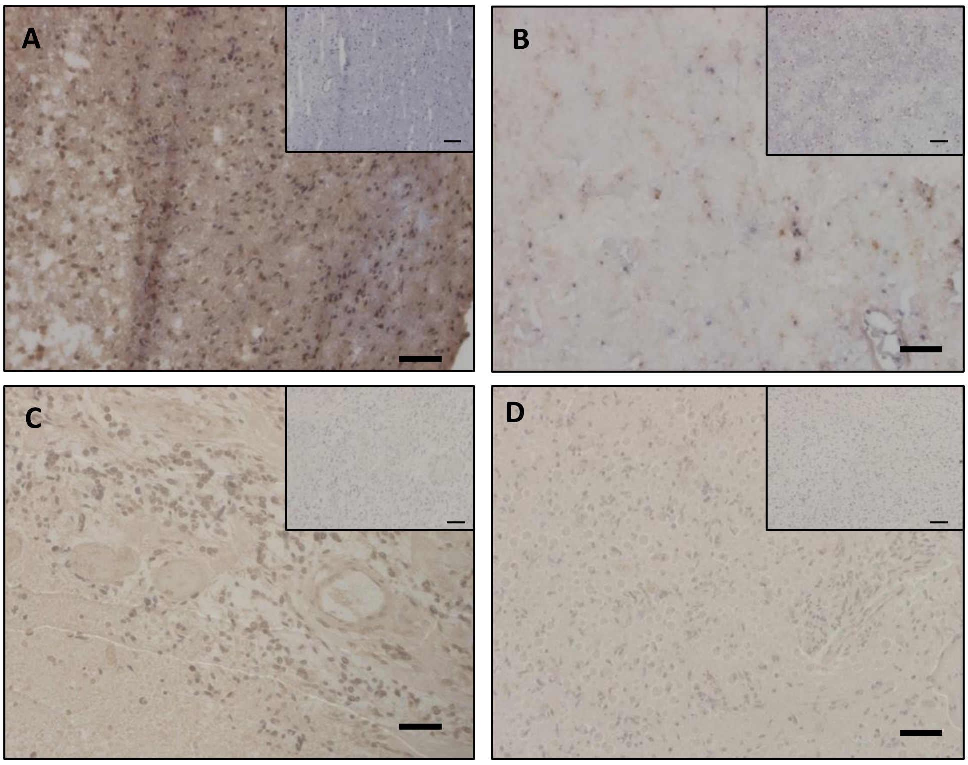


**Supplementary Figure 2. Immunohistochemistry showing high-level GD2 expression in glioblastoma (GBM) and H3-K27M diffuse intrinsic pontine glioma (DIPG) tumor tissue but not normal brain or H3-wildtype DIPG.** Tissue specimens from GBM (n = 9), DIPG (n=4) or non-involved brain tissue (n = 2) sections were stained by immunohistochemistry using anti-GD2 mAb (clone 14g2a) as primary antibody. Negative controls used an irrelevant IgG2a (insets). Scale bar = 50mm. (**A**) Representative GBM. **(B**) Non-involved brain tissue. (**C**) Representative H3-K27M mutation DIPG. (**D**) Representative H3-wildtype DIPG.

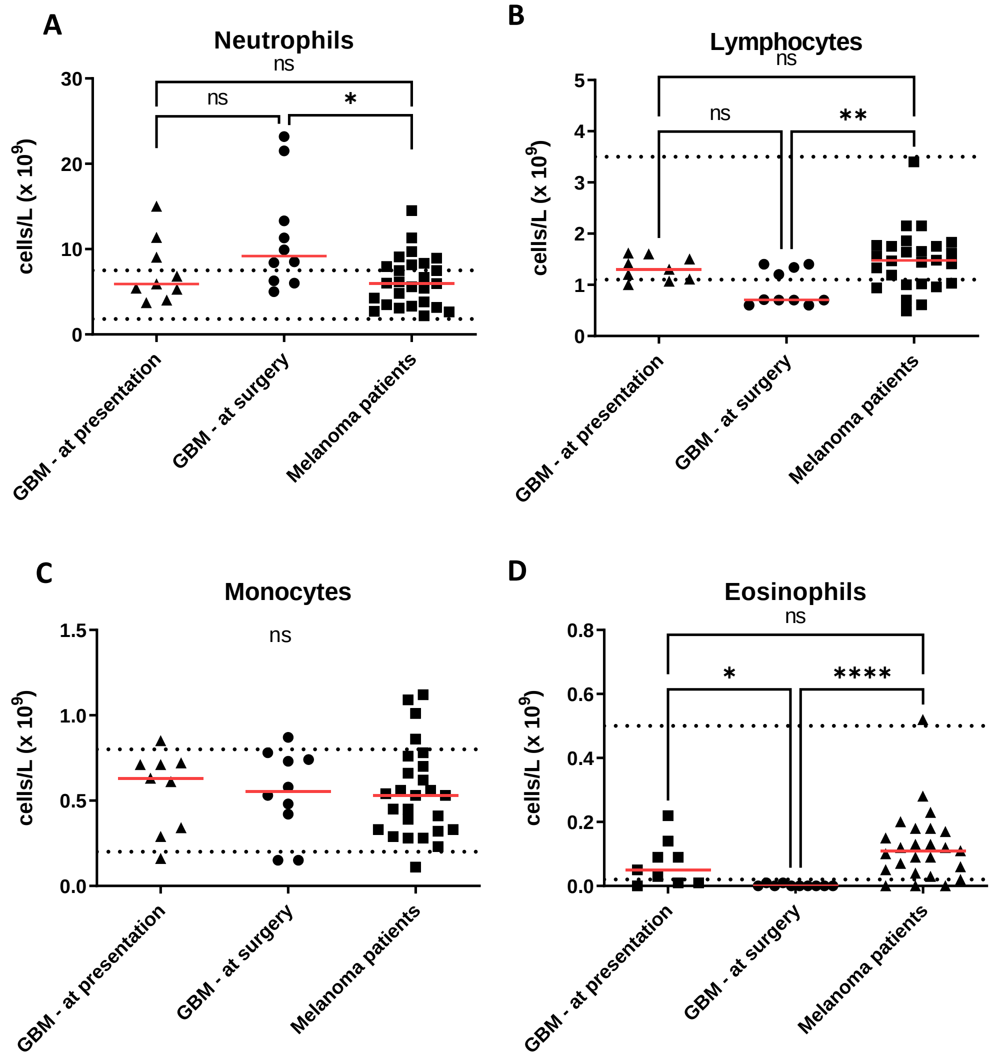
A
Neutrophils
ns
ns
*
cells/L (x 10^9)
30
20
10
0
GBM - at presentation
GBM - at surgery
Melanoma patients
B
Lymphocytes
ns
ns
**
5
4
3
2
1
0
C
Monocytes
ns
1.5
1.0
0.5
0.0
D
Eosinophils
ns
*
****
0.8
0.6
0.4
0.2
0.0

E

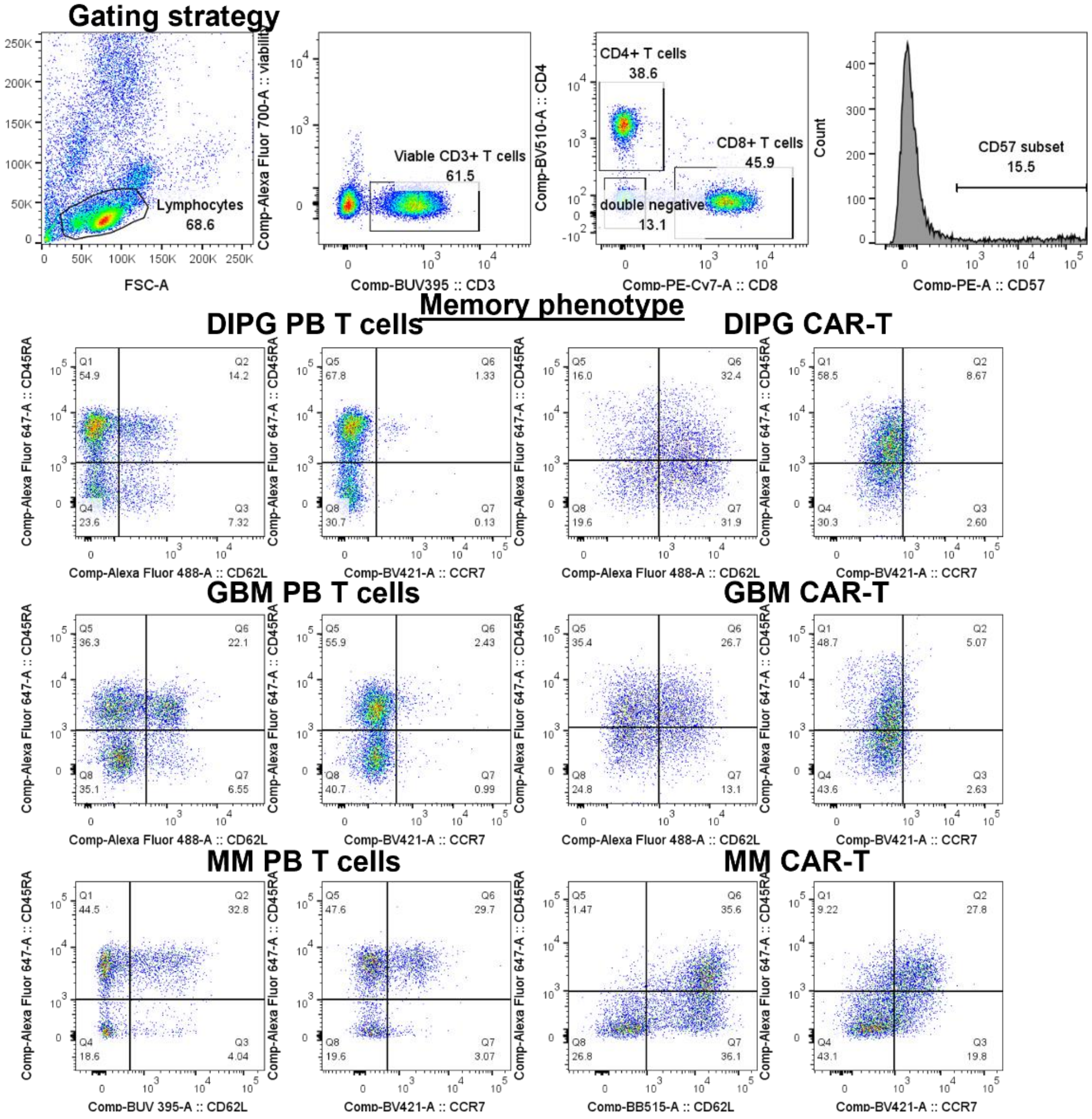


**Supplementary Figure 3. Blood leukocyte changes in GBM.** Patient pathology reports were used to determine numbers of **A)** neutrophils, **B)** lymphocytes, **C)** monocytes and **D)** eosinophils for GBM patients at presentation and again at surgery. A selection of melanoma patients is provided for comparison. Dotted lines indicate healthy normal range. **E)** Representative histograms showing gating strategy and patterns of CCR7 and CD62L expression for peripheral blood (PB) T cells and CAR-T cells from GBM, DIPG and metastatic melanoma (MM) patients.

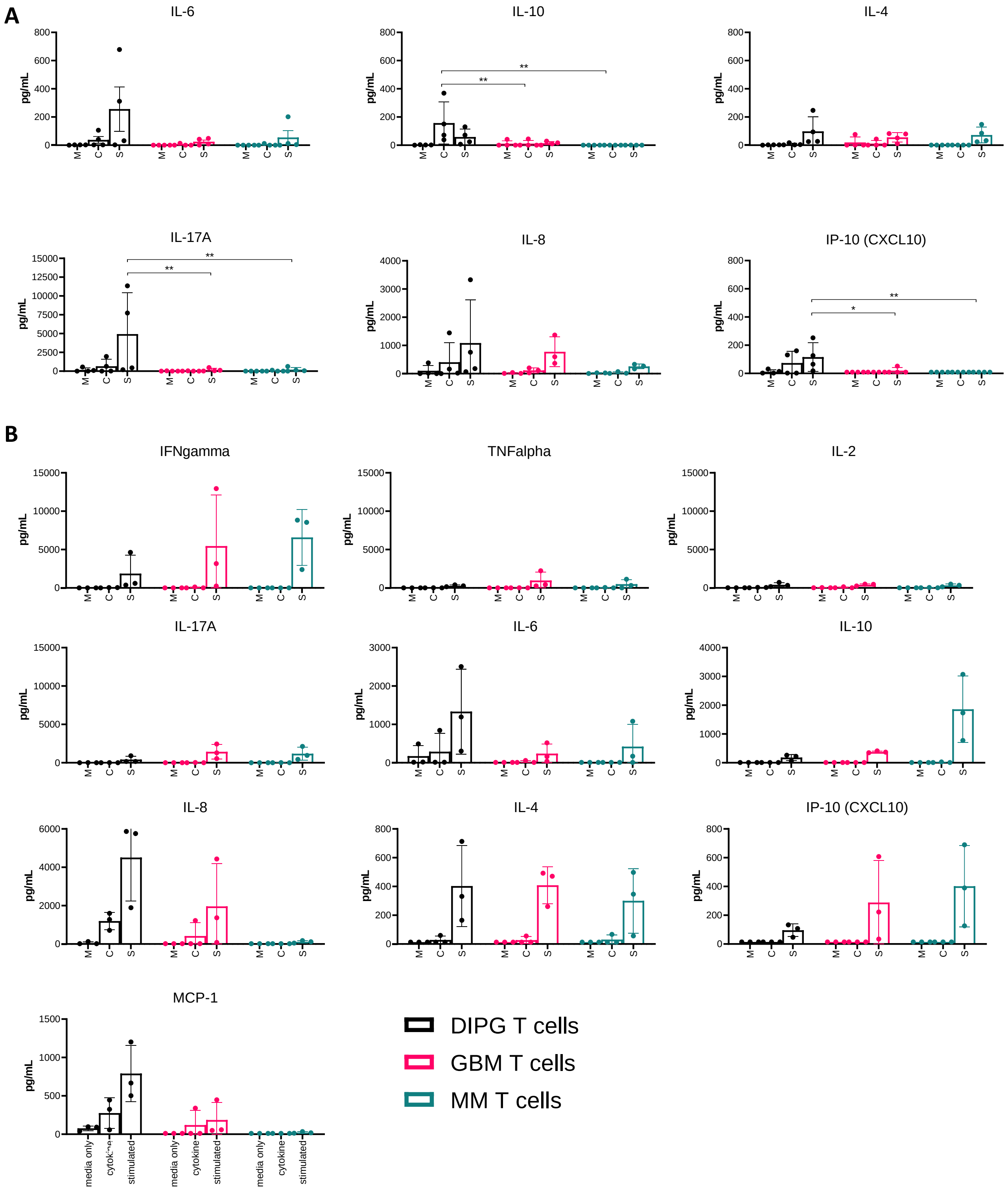


**Supplementary Figure 4. Cytokine production from CAR-T cell and peripheral blood T cells sorted from GBM and DIPG patients.** T cells were cultured with media (M) only, media plus IL-7 and IL-15 cytokine (C) for homeostatic proliferation, or media and plate-bound anti-CD3 and anti-CD28 antibody (S) (1ug/mL) for stimulation for 72 hours. **A)** Cytokines produced from CAR-T cells ]. **B)** Cytokines produced from peripheral T cells. T cells were sorted from GBM, DIPG and melanoma patients' blood using pan-T cell selection magnetic beads. This negative selection method allows T cells to remain untouched.

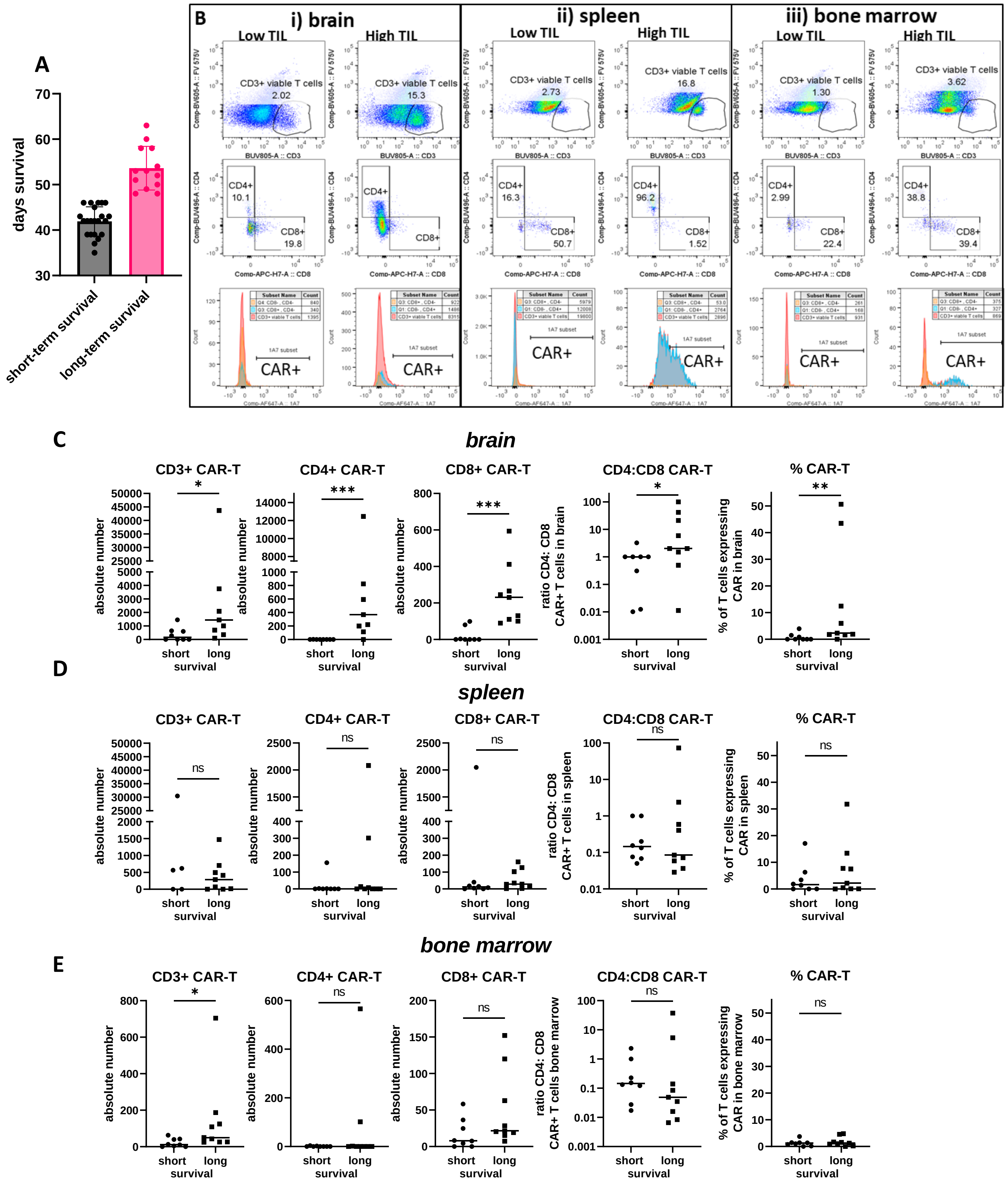
A
days survival
short-term survival
long-term survival
B
i) brain
ii) spleen
iii) bone marrow
Low TIL
High TIL
CD3+ viable T cells
CD4+
CD8+
CAR+
C
brain
CD3+ CAR-T
CD4+ CAR-T
CD8+ CAR-T
CD4:CD8 CAR-T
% CAR-T
absolute number
ratio CD4: CD8 CAR+ T cells in brain
% of T cells expressing CAR in brain
short long survival
D
spleen
ratio CD4: CD8 CAR+ T cells in spleen
% of T cells expressing CAR in spleen
E
bone marrow
ratio CD4: CD8 CAR+ T cells bone marrow
% of T cells expressing CAR in bone marrow

**Supplementary Figure 5. Flow cytometry CAR-T cell enumeration and statistical analysis for mice treated with CAR-T cells from healthy or GBM donors CAR-T cells. A)** Short and long term survival groups. Long term survivors defined as surviving longer than 46 days. **B)** Representative dot plots and histogram of CD3+, CD4+ and CD8+ CAR-T cells in **i)** brain **ii)** spleen and **iii)** bone marrow. Examples of mice with low and high numbers of tumor-infiltrating lymphocytes (TIL) of T-cell type were selected. The low-TIL mouse had a survival of 43 days, the high-TIL mouse had a survival of 63 days. Both mice were treated with cells from the same donor (GBM patient BT 48). **C-E)** Statistical analysis of association between CAR-T absolute numbers, CD4: CD8 CAR-T cell ratio and percentage of human $CD3^{+}$ T cells expressing surface CAR and groups of mice split according to survival time.

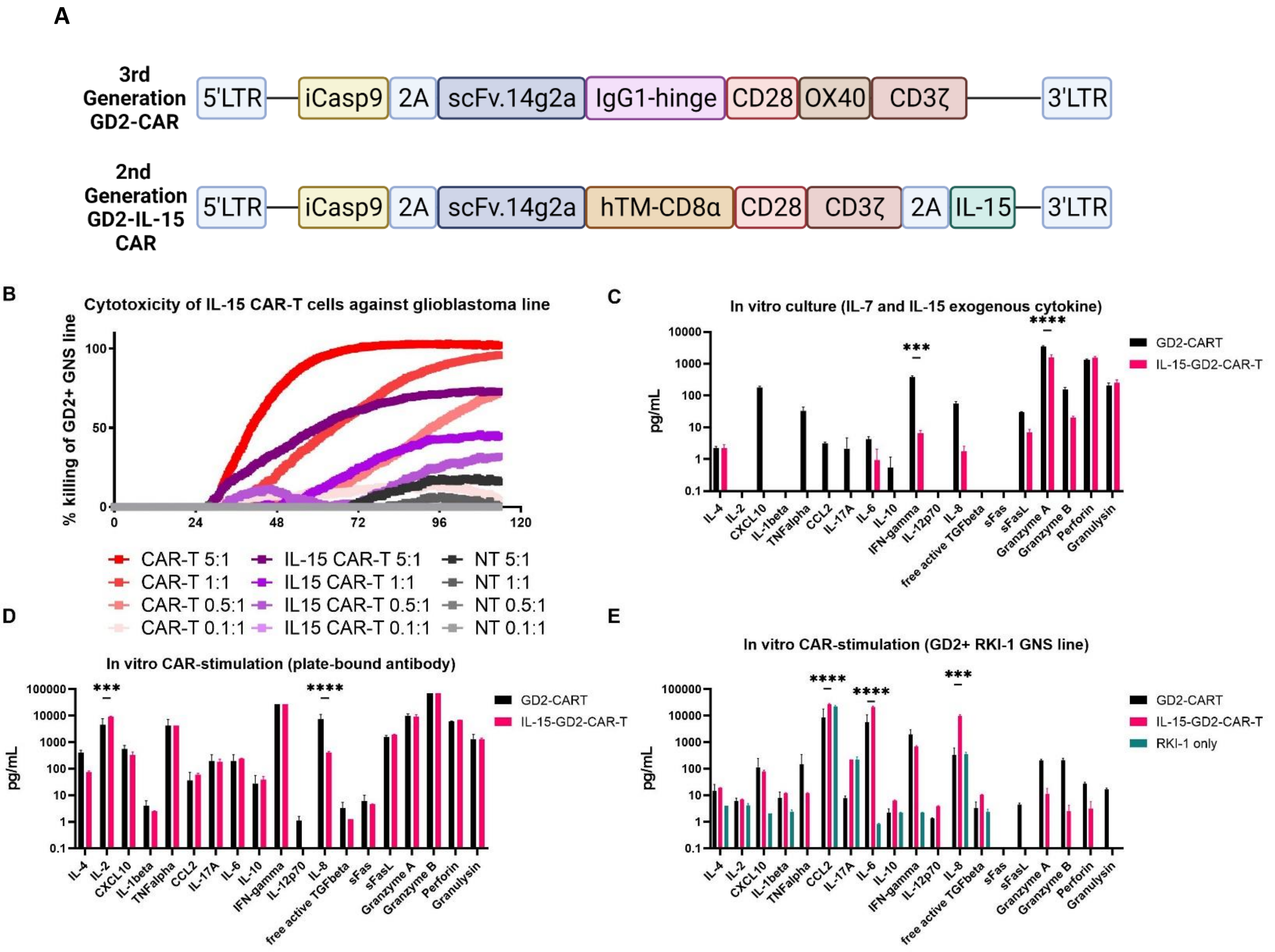


**Supplementary Figure 6**. **IL-15 transgene effects on in vitro functionality. A)** Com parison of the two clinical retrovectors. The second-generation GD2-CAR-IL-15 construct is characterised by a slightly shorter $V_H$ sequence, a longer 20aa linker between the $V_L$ and $V_H$ domains, and hinge and transmembrane domains from CD8alpha (as opposed to the short IgG1 hinge and CD28 transmembrane domain used in the third-generation GD2-CAR construct), and including the IL-15 transgene, as described in Chen et al. Clin Cancer Res 2019. **B)** In vitro cytotoxicity as assessed by co-culture with a GD2$^+$ GNS line (RKI-1) in a real-time impedance-based cytotoxicity assay. Healthy donor GD2-CAR-IL-15 T cells (purple lines) were compared to healthy donor third-generation GD2-specific CAR-T cells (red lines) at 5:1, 1:1, 0.5:1 and 0.1:1 effector to target ratios. Cytokine bead arrays (Legendplex) were performed using supernatants from cultured healthy donor GD2-CAR-IL-15 T cells were compared to healthy donor third-generation GD2-specific CAR-T cells after 72 hours with **C)** exogenous cytokines (IL-15, 5ng/mL and IL-7, 10ng/mL) or **D)** stimulation with plate-bound 1A7 anti-CAR antibody (2μg/mL). **E)** Supernatants from the real-time cytotoxicity assay were also harvested and tested for cytokines.

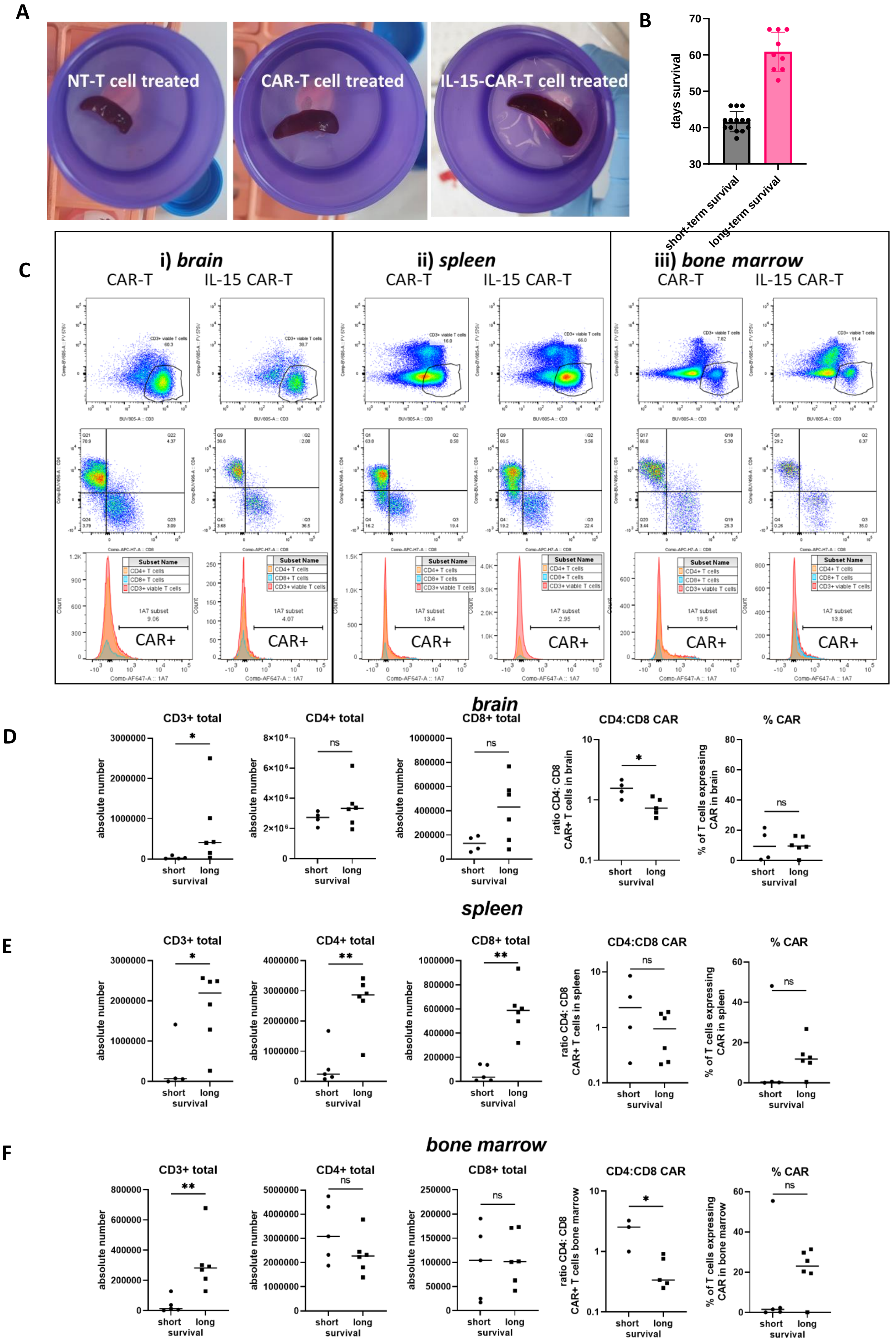
A
NT-T cell treated
CAR-T cell treated
IL-15-CAR-T cell treated
B
days survival
short-term survival
long-term survival
C
i) brain
ii) spleen
iii) bone marrow
CAR-T
IL-15 CAR-T
CAR+
D
brain
CD3+ total
CD4+ total
CD8+ total
CD4:CD8 CAR
% CAR
absolute number
ratio CD4: CD8 CAR+ T cells in brain
% of T cells expressing CAR in brain
short long survival
E
spleen
ratio CD4: CD8 CAR+ T cells in spleen
% of T cells expressing CAR in spleen
F
bone marrow
ratio CD4: CD8 CAR+ T cells bone marrow
% of T cells expressing CAR in bone marrow

**Supplementary Figure 7**. **IL-15 transgene effects on CAR-T cell engraftment, infiltration and mouse survival. A)** examples of increased spleen size for GD2-CAR-IL-15 T cell treated mice. **B)** Short- and long-term survival groups. Long-term survivors defined as surviving longer than 46 days. **C)** Representative dot plots and histograms of $CD3^+$, $CD4^+$ and $CD8^+$ CAR-T cells in **i)** brain, **ii)** spleen, and **iii)** bone marrow. Statistical analysis of CAR-T absolute numbers, CD4: CD8 CAR-T cell ratio and percentage of human $CD3^+$ T cells expressing surface CAR in **D)** brain, **E)** spleen, and **F)** bone marrow and association with mouse survival.

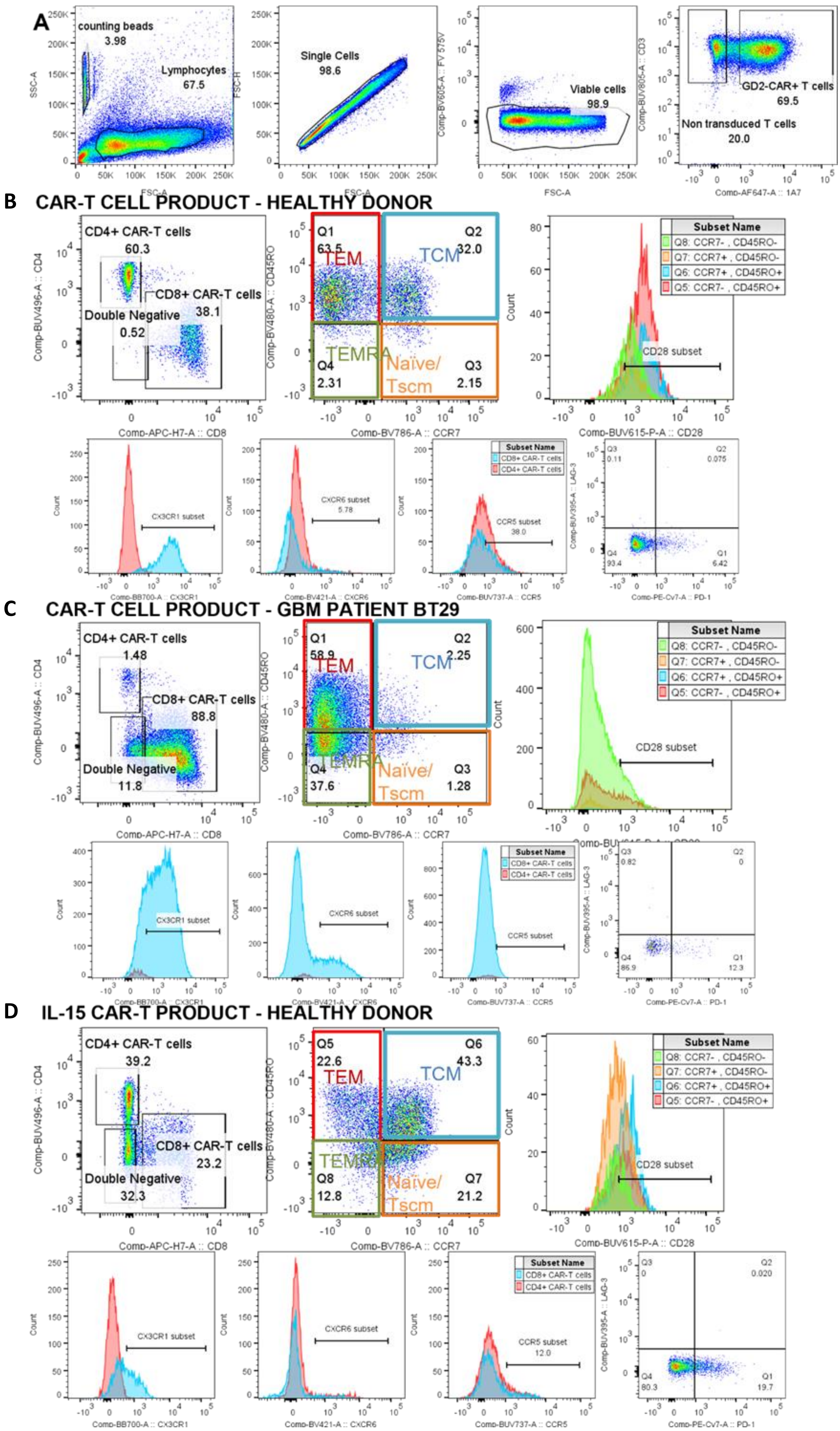

A
counting beads
3.98
Lymphocytes
67.5
Single Cells
98.6
Viable cells
98.9
GD2-CAR+ T cells
69.5
Non transduced T cells
20.0
B CAR-T CELL PRODUCT - HEALTHY DONOR
CD4+ CAR-T cells
60.3
CD8+ CAR-T cells
38.1
Double Negative
0.52
TEM
TCM
TEMRA
Naïve/ Tscm
Subset Name
Q8: CCR7- , CD45RO-
Q7: CCR7+ , CD45RO-
Q6: CCR7+ , CD45RO+
Q5: CCR7- , CD45RO+
CD28 subset
C CAR-T CELL PRODUCT - GBM PATIENT BT29
CD4+ CAR-T cells
1.48
CD8+ CAR-T cells
88.8
Double Negative
11.8
D IL-15 CAR-T PRODUCT - HEALTHY DONOR
CD4+ CAR-T cells
39.2
CD8+ CAR-T cells
23.2
Double Negative
32.3

**Supplementary Figure 8**. **Gating strategy and example histograms for 14-color flow cytometric analysis. A)** Gating strategy to detect CAR-T cells. Representative dot plots to determine phenotype for the GD2-specific third-generation CAR-T cell product produced from **B)** healthy donor, **C)** GBM donor, and **D)** healthy donor GD2-specific second-generation CAR-T cells expressing IL-15.

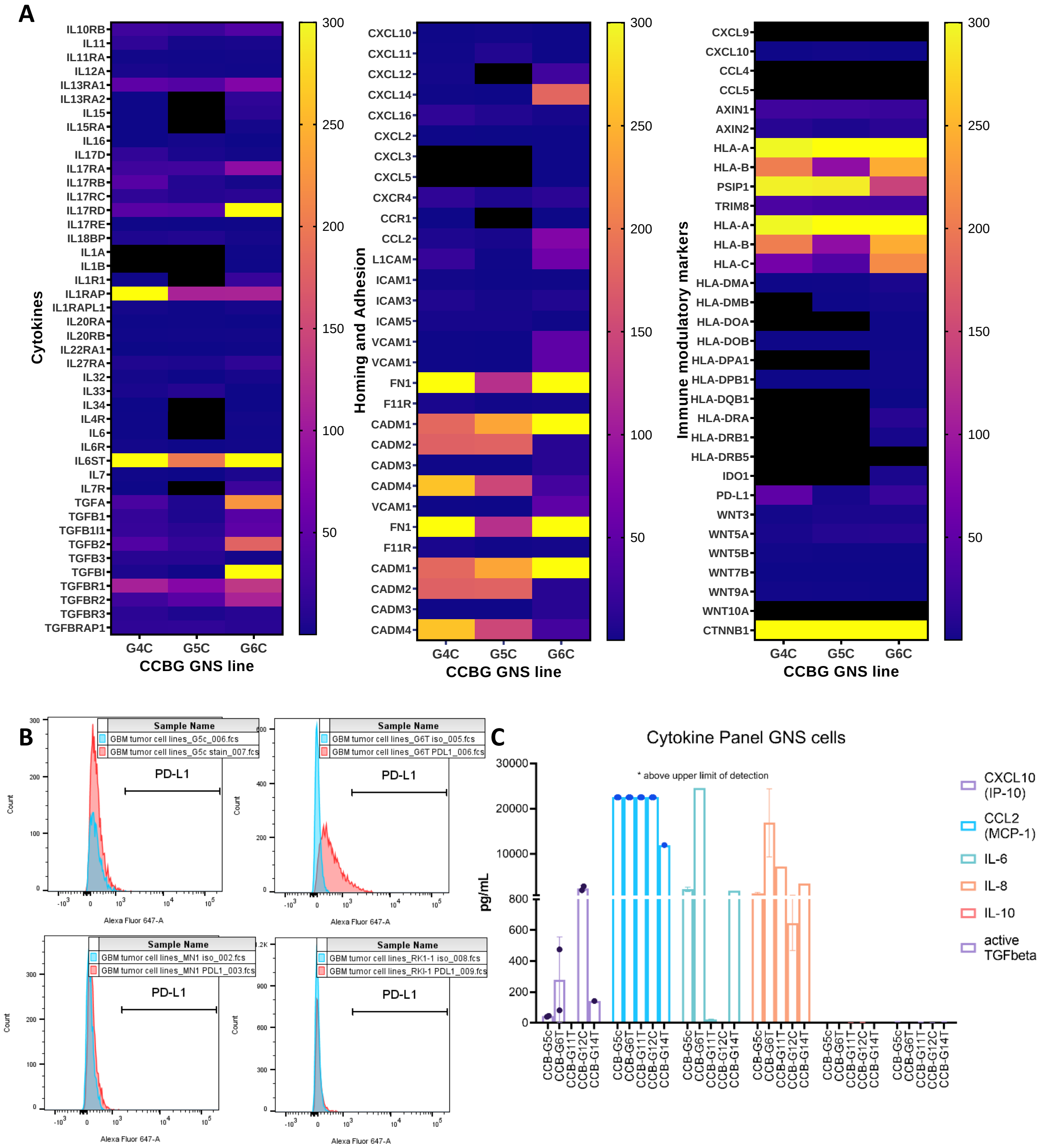


**Supplementary Figure 9. RNA, flow cytometric and immunofluorescence microscopy analysis of GBM microenvironmental factors that may affect CAR-T cell function. A)** Bulk RNA sequencing was performed on 3 GNS lines (CCB-G4C, **CCB-G5C**, CCB-G6C), sequenced in triplicate). Heat maps showing expression of known immune-modulatory genes were then produced. **B)** Flow cytometric analysis of 4 GNS lines for PD-L1 (**CCB-G5C,** CCB-G6T, MN1, RK-1). **C)** Cytometric bead array analysis of secreted cytokines and chemokines from 5 GNS lines (**CCB-G5C,** CCB-G6T, CCB-G11T, CCB-G12C, CCB-G14T). CCB-G5C is the line used in the xenograft model.
