## Supplementary Methods for "GD2-targeting CAR-T cells enhanced by transgenic IL-15 expression are an effective and clinically feasible therapy for glioblastoma"

**RNA sequencing and analysis**

Cryopreserved dissociated mouse brains hosting human glioblastoma tumors were rapidly thawed and Trizol added before RNA extraction was performed by standard techniques. Dual replicate polyA$^+$ enriched RNA-seq libraries from 8 biological samples for each of 4 treatments (untreated, NT-T treated, GD2-CAR-T treated, and GD2-CAR-IL-15-T treated) were multiplexed and sequenced on two separate Illumina NextSeq 500 runs using the stranded, single end protocol with a read length of 75.

Raw reads were adaptor trimmed and filtered for short sequences using cutadapt v2.10 setting minimum-length option to 18, error-rate 0.2 and overlap 5. The resulting FASTQ files averaging 30 million reads per sample were analysed and quality checked using the FastQC program (http://www.bioinformatics.babraham.ac.uk/projects/fastqc). Reads were mapped against both the mouse (mm10) and the human reference genome (hg38) using the STAR spliced alignment algorithm (version 2.7.7a with default parameters and --chimSegmentMin 20, --quantMode GeneCounts) in a two-stage iterative process. The first round was used to align and remove duplicate reads using UMI-tools (version 1.0.0 with default parameters). The resulting deduplicate data was realigned returning an average unique alignment rate of 53% against the mouse and 34% against the human reference genomes. Differential expression analysis was evaluated from TMM normalized gene counts using R (version 4.1.3) and edgeR (version 3.34.1). Alignments were visualised and interrogated using the Integrative Genomics Viewer v2.3.80. Graphical representations of differentially expressed genes were generated using Glimma.

**Human and mouse tissue histology and Immunofluorescence**

For Hematoxylin and eosin staining the frozen tissue sections were fixed with cold 10% neutral buffered formalin for 10 minutes, then briefly washed with tap water. Mayers haematoxylin (Agilent Australia; Mulgrave, Victoria, Cat# S330930-2) was used to stain the nuclei and blued by in-house prepared Scott's tap water (2% $MgSO_4$ solution buffered with $NaHCO_3$). Sections were then thoroughly rinsed in water and 95% EtOH before counterstained with an alcoholic solution of eosin (Sigma-Aldrich Australia; North Ryde, NSW cat# HT110116) for three minutes. Following this, sections were dehydrated through several baths of ethanol and cleared with xylene twice before mounted in DPX mountant (Sigma-Aldrich).

Nanozoomer captured whole tissue images were provided to independent pathologists (Gribbles, Australia) to asses as digital slides in a blinded fashion. The human pathologist scored the tumor for cellularity, vascular proliferation and mitotic features. The veterinary pathologist scored the adjacent brain for vascular proliferation, gliosis and perivascular cuffing.

For GD2 staining of human GBM and adjacent normal brain sections, tissues were fixed for 20 minutes in Cytofix/Cytoperm (BD) at room temperature (RT), washed in PBS and blocked using 10% human plasma in PBS containing 1% bovine serum albumin (BSA) for 30 minutes at RT. Sections were incubated with mouse anti-GD2-AF647 (14.G2A; BD) or isotype control mouse IgG2a-AF647 (G155-178; BD) at dilutions of 1:100 for 12 hours at 4°C. After washing, sections were mounted using ProLong Gold anti-fade mounting medium with DAPI (Life Technologies), cured overnight, and sealed with nail varnish before viewing.

For staining of mouse brain sections, contiguous sections were fixed as described above and blocked using 10% normal rat serum and 10% normal goat serum in PBS containing 1% BSA for 30 minutes at RT. Tissue sections were stained using CD3 and CD31 or CD3 and GD2 antibodies. For CD3 and CD31 staining, 5µg/mL purified mouse anti-CD3 (UCHT1; BioLegend) and 4µg/mL purified hamster anti-CD31 (2H8; Thermo Fisher Scientific) were incubated with tissue sections for 12 hours at 4°C. Following washing, tissue sections were incubated with goat anti-mouse IgG AF647 (Thermo Fisher Scientific) and goat anti-hamster IgG AF488 (Abcam) detection antibodies at a dilution of 1:500 for 1 hour at RT. For CD3 and GD2 staining, 5µg/mL purified mouse anti-CD3 alone was incubated with tissue sections for 12 hours at 4°C. Following washing, tissue sections were incubated with goat anti-mouse IgG AF647 at a dilution of 1:500 for 1 hour at RT. Following washing, sections were blocked using 10% normal mouse serum in PBS containing 1% BSA for 30 minutes at RT. Sections were then incubated with mouse anti-GD2-FITC (14.G2A; BD) at a dilution of 1:100 for 2 hours at RT. Following final antibody staining, sections were washed and mounted for viewing as described above.

Whole-slide imaging was performed on a Zeiss Axio Scan.Z1 slide-scanner using 20x objective and ZEN 3.1 Blue system software. Fluorescence overlays were created by merging channels and applying false colour using FIJI (ImageJ, National Institutes of Health). ROI were generated mapped to the outline of tissue sections and mean intensity measured for each channel. Counting of CD31+ vessels (identified as rings of CD31+ cells surrounding a black lumen) across an entire section was performed manually. As this was deemed the most subjective analysis it was performed blinded manner with mouse treatment groups de-identified. All other analyses were not blinded.

**Flow Cytometry**

Thawed cells were rested for 2 hours and then stained with conjugated antibodies for 30 minutes on ice in Brilliant Stain Buffer (BSB; Becton Dickinson). Fixable viability stains were performed for 10 minutes in PBS, followed by washing and fixation (4% paraformaldehyde). Antibodies for multi-color panels are listed in **Supplementary Table 2**. Stained cells were analyzed on a BD LSR Fortessa (BD Biosciences) or a BD FACSymphony A5 (BD Biosciences) with FlowJo Software V 10.8.1 (BD Biosciences). Fluorescence minus one (FMO) controls were used to establish negative gates for peripheral blood and cultured CAR-T cells, whereas fully stained, dissociated, tissues from control mice were used to establish gates for *ex vivo* CAR-T cell analysis. After staining cells were transferred to TruCount tubes (BD Biosciences) to enable absolute cell numbers to eb determined. Cytokines were analysed using the Legendplex Cytometric Bead Array (Human Essential Immune Response kit). Cells were stimulated with plate-bound 2µg/mL 1A7 (to stimulate the CAR) or cultured in media +/- IL-7 and IL-15 for 3 days and then cells and supernatants were collected for analysis by flow cytometry.

**Real Time Cell Adhesion Cytotoxicity Assay**

Cytotoxicity assays were performed using the Maestro Z impedance assay (Axion Biosystems). For this assay, 10,000 $GD2^+$ LAN-1 or 12,000 GNS or DIPG tumor cells were plated in a CytoView-Z 96-well plate and cultured for 16 hours before the addition of CAR-T cells at ratios of 0.1:1, 0.5:1, 1:1, 5:1 and cultured for an additional 120 hours. Impedance data was normalized to the time-point just before addition of CAR T cells (i.e., impedance at time 0 = 1). Analysis of cytotoxicity and time to 50% killing (KT50) was performed using the Axis Z software (Axion Biosystems) using target cells cultured alone as a baseline.
